# Structure and misfolding of the flexible tripartite coiled coil domain of glaucoma-associated myocilin

**DOI:** 10.1101/154112

**Authors:** Shannon E. Hill, Elaine Nguyen, Rebecca K. Donegan, Anthony Hazel, James C. Gumbart, Raquel L. Lieberman

## Abstract

Glaucoma-associated myocilin is a member of the olfactomedins, a protein family broadly involved in neuronal development and human disease. Molecular studies of the myocilin N-terminal coiled coil demonstrate a unique tripartite architecture: a disulfide-linked, parallel dimer-of-dimers Y-shaped molecule, with distinct tetramer and dimer regions. The structure of the C-terminal 7-heptad dimer elucidates an unexpected repeat pattern involving electrostatic inter-strand stabilization. Molecular dynamics simulations reveal an alternate conformation in which the terminal inter-strand disulfide bond limits the extent of unfolding and results in a kinked configuration. Taken together, full-length myocilin is also branched, with two pairs of C-terminal olfactomedin domains. Selected variants within the N-terminal region alter the apparent quaternary structure of myocilin but do so without compromising stability or causing aggregation. In addition to increasing our structural knowledge of extracellular coiled coils for protein design and biomedically important olfactomedins, this work broadens the scope of protein misfolding in the pathogenesis of myocilin-associated glaucoma.

**Highlights:**

- Glaucoma-causing extracellular protein associated with amyloid forming propensity
- Structural studies confirm tripartite parallel dimer-of-dimers coiled coil
- Leucine zipper exhibits non-canonical heptad repeat pattern with disulfide cap
- Glaucoma-associated variants in tetramer alter structure but not stability

**eTOC blurb:** Hill et al describe the structure of the coiled-coil region of myocilin, the extracellular olfactomedin family member closely associated with the ocular disorder glaucoma. Myocilin’s coiled coil adopts a unique Y-shaped parallel dimer-of-timers employing an unusual heptad repeat pattern. Selected disease variants alter quaternary structure.

## Introduction

Olfactomedins (PFAM: PF02191), a family of extracellular, modular, multidomain proteins each with a 5-bladed β-propeller olfactomedin (OLF) (Donegan et al., 2015) domain, are broadly involved in eukaryotic development (Tomarev and Nakaya, 2009; Zeng et al., 2005). The majority of olfactomedin subfamilies also contain a high probability signal sequence for cellular secretion followed by homo-oligomerizing coiled-coil (CC) domains of varying lengths (Figure 1). Despite their prevalence in multicellular organisms and implication in a variety and increasing number of human diseases (Anholt, 2014), however, the molecular architecture and specific functions of olfactomedins remain largely unknown.

**Figure 1.**
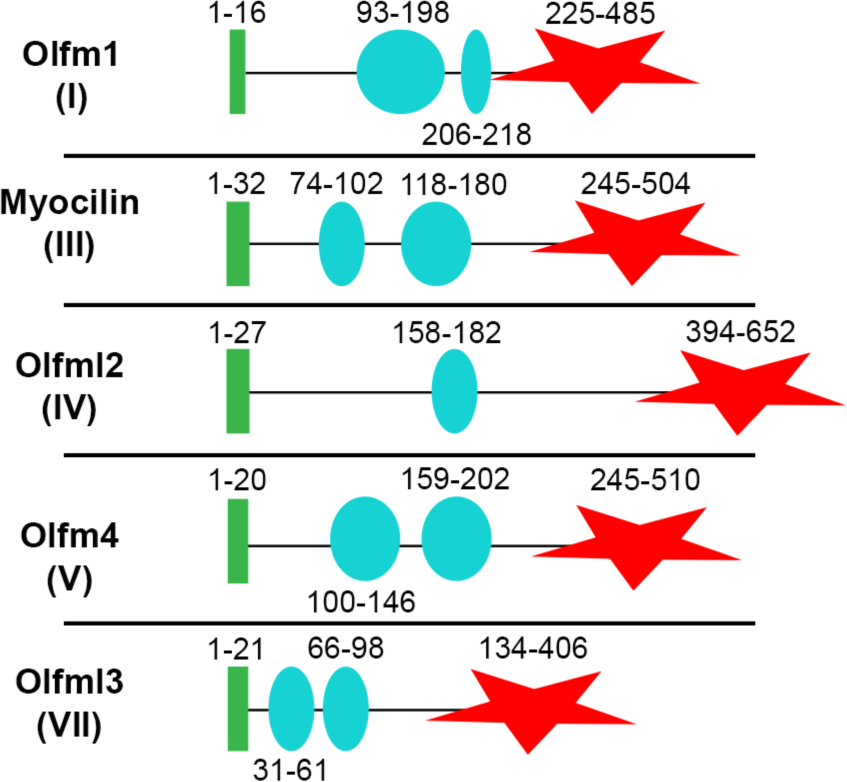
Distinctive domain structures among CC-containing olfactomedin subfamilies. Green bar, predicted signal sequence for cellular secretion; cyan circles, predicted CC regions; red star, olfactomedin domain. Subfamily II is membrane-anchored; subfamily VI has a collagen domain.

The structures of CCs, one of the most common structural domains in nature (Wolf et al., 1997) and of considerable interest for protein design and biomaterials (Lupas and Bassler, 2017; Woolfson, 2005), can be challenging to predict *in silico*, particularly beyond dimers that are common among extracellular CCs (Kammerer, 1997). The CCs encoded within olfactomedins tend to have non-canonical heptad repeat (*abcdefg*)_n_ patterns, diverging from the regular period of hydrophobic residues at positions ‘*a*’ and ‘*d*’ and electrostatic residues at positions ‘*e*’ and ‘*g*’ (Mason and Arndt, 2004). Thus, bioinformatics programs detect CCs in olfactomedins, but vary in predictions for register, stoichiometry, and orientation. Olfactomedin-1 (subfamily 1) is the only olfactomedin protein for which a supramolecular organization has been experimentally characterized: an apparent V-shaped molecule seen by transmission electron microscopic imaging and small angle X-ray scattering (SAXS) envelope of the full-length protein(Pronker et al., 2015). However, because olfactomedin family CC domains share very low sequence similarity (< 15% identity overall, Supplementary Figure S1), results from olfactomedin-1 are not predictive of other olfactomedin family members.

Here we characterized the unique architecture of full-length myocilin (olfactomedin subfamily 3), one of the best-studied olfactomedins, in molecular detail. Myocilin is expressed at high levels in the trabecular meshwork (TM), an extracellular matrix in the anterior segment of the eye that forms part of the anatomical ultrafiltration system for aqueous humor (Abu-Hassan et al., 2014), and is a critical tissue for maintaining intraocular pressure. The myocilin CC-containing region has been reported to interact with matrix components (Filla et al., 2002; Ueda et al., 2002; Wentz-Hunter et al., 2004) and confer its adhesion properties (Goldwich et al., 2009; Wentz-Hunter et al., 2004), but there is much still to learn about the structure and biological function of myocilin. Myocilin is associated with two subtypes of glaucoma (Polansky et al., 1997; Rozsa et al., 1998; Stone et al., 1997), a leading cause of blindness worldwide typically associated with elevated intraocular pressure (Quigley and Broman, 2006). Mutations in myocilin are causative for Mendelian-inherited form of open angle glaucoma (~4% of ~70 million), the most common subtype (Hewitt et al., 2006). OLF-resident myocilin variants are destabilized (Burns et al., 2010; Burns et al., 2011) and prone to amyloid-like aggregation (Hill et al., 2014; Orwig et al., 2012), which is believed to cause a toxic gain of function, leading to early-onset glaucoma (Stothert et al., 2016).

We present biophysical and biochemical experiments demonstrating that the myocilin CC domain is a distinct Y-shaped molecular entity composed of two parallel dimer-of-dimers that are stabilized on the three ends by disulfide bonds. The 1.9 Å resolution X-ray structure of the C-terminal 7 heptads of the CC further reveals an unexpected repeat pattern involving inter-strand stabilization between oppositely charged residues at heptad position ‘*a*’ and ‘*g*’. Molecular dynamics simulations demonstrate that the C-terminal capping disulfide bond limits unfolding of the CC and enables an alternative kinked structure to form. Selected glaucoma-associated mutations within the N-terminal coiled coil region adopt non-native structures but are not destabilized or aggregation prone. Our study increases structural knowledge of extracellular coiled coils, biomedically important olfactomedins, and broadens our appreciation of protein misfolding in the pathogenesis of myocilin-associated glaucoma.

## Results and Discussion

### The myocilin coiled-coil imparts a unique tri-branched supramolecular architecture

After unsuccessful attempts to express soluble full-length myocilin in *E. coli* or purify sufficient quantities from spent media of primary human TM cells (not shown), we generated well-behaved constructs of myocilin outside of the OLF domain whose structure we solved previously (Donegan et al., 2015). Biophysical characterization of NTD_33-227,_ which includes linker regions before and after the predicted CC domain (Figure 1), reveals the typical circular dichroism (CD) signatures for α-helices. Thermal unfolding is reversible (Figure 2a, Table 1, melting temperature ((Tm) ~ 67 °C), a value considerably higher than the ~52 °C measured for the OLF domain (Orwig and Lieberman, 2011). To investigate the contribution of Cys residues to the stability and oligomeric state of NTD_33-227,_ Cys-to-Ser mutants were prepared. The Cys residues thermally stabilize NTD_33-227_ nearly 20 °C, with Cys185 contributing the biggest thermal increase (Table 1), but do not change the elution profile from size exclusion chromatography (SEC) and thus the apparent quaternary structure (Figure 2b). In each case, the predominant species present upon disulfide bond formation is a dimer (Figure 2c).

**Figure 2.**
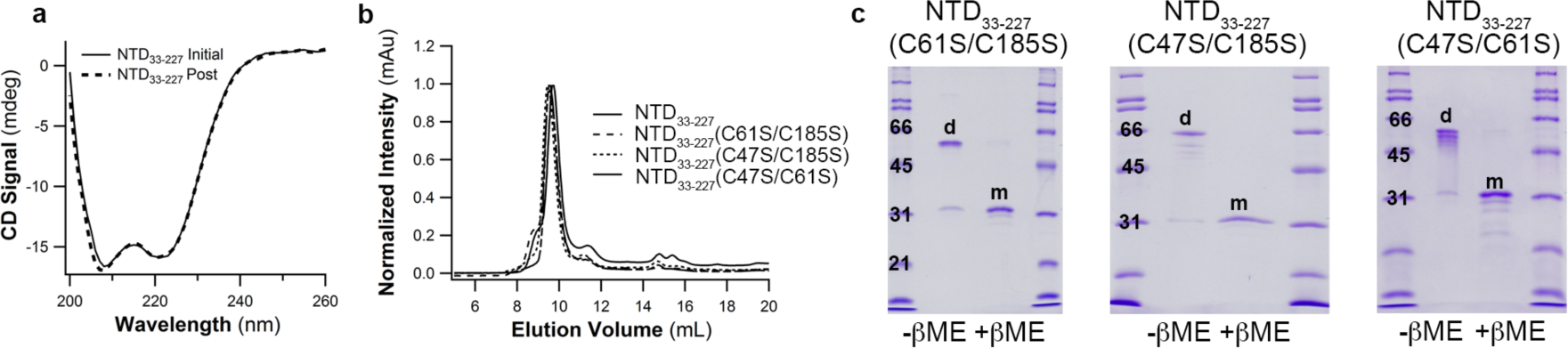
Biophysical characterization of NTD_33-227._ (a) Initial NTD_33-227_ circular dichroism spectrum reveals α-helical signature. Overlay with post-melt NTD_33-227_ spectrum confirms thermal unfolding is reversible. (b) SEC elution profiles demonstrate unchanged quaternary structure for wild type and Cys-to-Ser NTD_33-227_ variants. (c) SDS-PAGE analysis of NTD_33-227_ Cys-to-Ser variants under non-reducing conditions show predominantly dimer (d) species. Calculated molecular weight for NTD_33-227_ monomer (m) = 27 kDa.

**Table 1.**
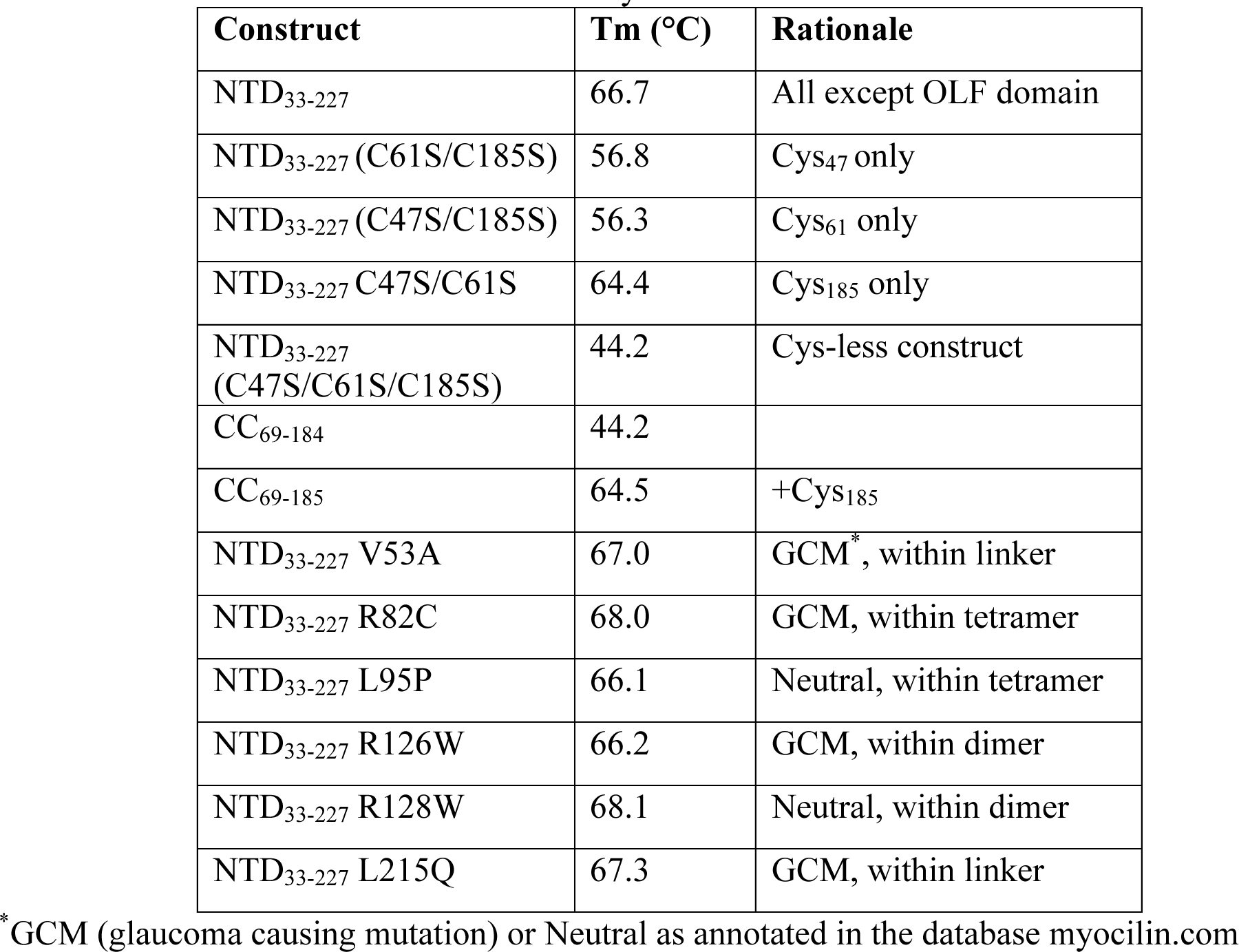
Tm values for constructs in this study.

To clarify further the oligomeric state and shape of the structural coiled coil of myocilin, we next prepared a CC construct bracketed by the two Cys residues closest to the predicted CC-(CC_60-185_). The SEC elution profile of this species is consistent with a 4- or 5-mer (Supplemental Figure S2a). The molecular envelope determined by SEC-SAXS analysis (Supplemental Figure S2) reveals a branched, tripartite structure that looks like the letter ‘Y’ (Figure 3a, Table 2). There is a short stem (3-4 nm wide x 4 nm long) and two arms (each 3-4 nm wide and 9-11 nm long) ~130° apart. The molecule appears nearly rotationally symmetric, but symmetry is broken due to apparent flexibility of the arms (Figure 3a).

**Figure 3.**
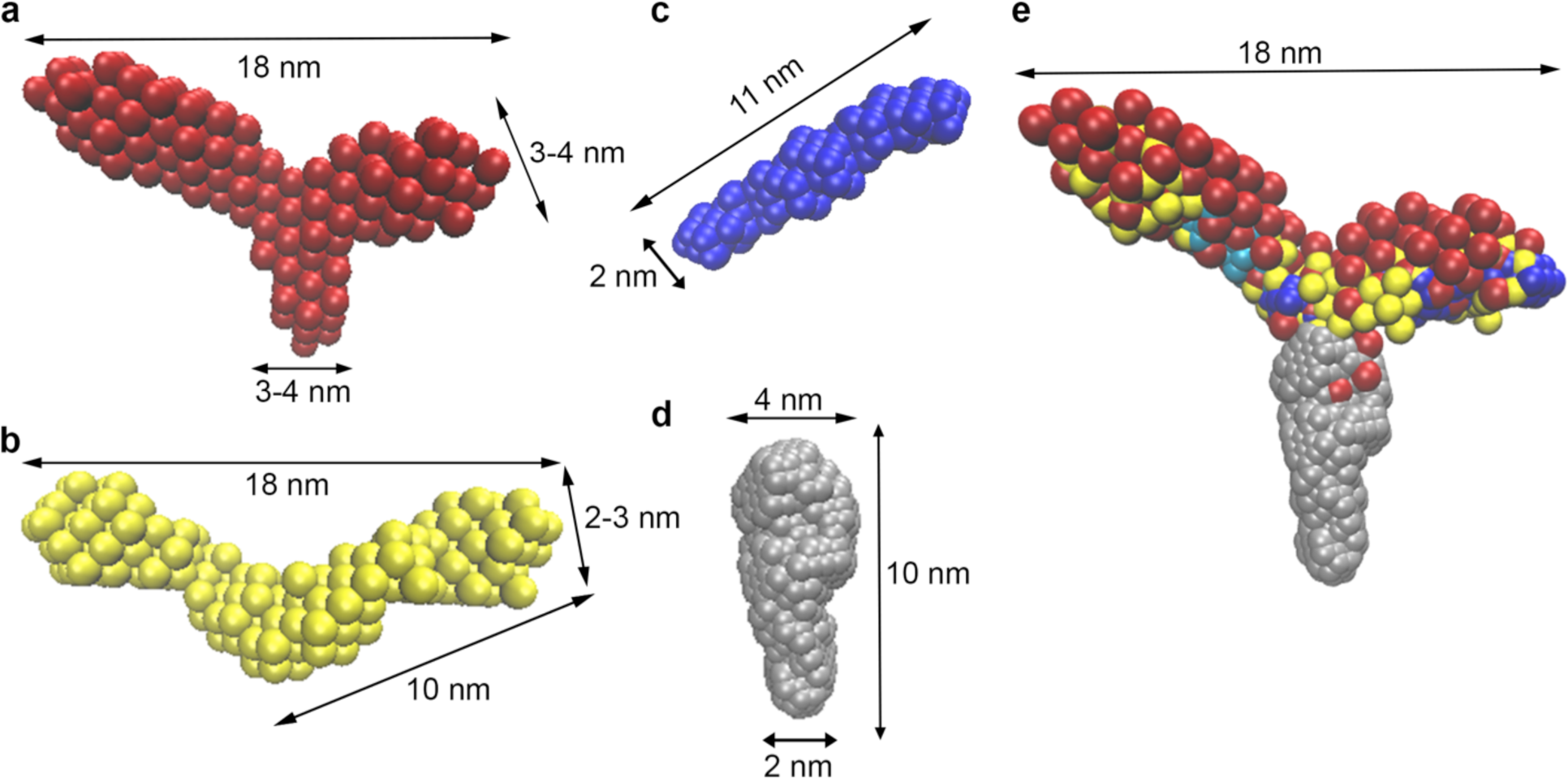
*Ab initio* models from SEC-SAXS reveal tripartite structure for myocilin CC. (a) CC_60-185_ is Y-shaped with arms spanning 18 nm and 130° apart from a stem ~3-4 nm in width. (b) CC_69-185_ is a P2-symmetric V-shaped molecule with similar dimensions arms of CC_60-185_. (c) CC_112-185_ is a P2-symmetric, ~11 nm long rod (d) CC_33-111_ is an oblong molecule with varying widths of 2-4 nm. (e) Overlay of SAXS molecular envelopes in a-d. See also Table 2.

**Table 2.** SEC-SAXS Statistics

| Construct | R<sub>g</sub><br>(Å) | Monomer<br>MW | SAXS Mol<br>(Oligomer) | Chi<sup>2</sup> | NSD |
| --- | --- | --- | --- | --- | --- |
| CC<sub>60-185</sub> | 60.2 | 14.6 kDa | 72 kDa (4.9) | 0.9692 | 0.900 |
| CC<sub>69-185</sub> | 56.5 | 13.7 kDa | 49 kDa (3.6) | 1.104 | 0.695 |
|  |  |  |  | 1.181 (P2) | 1.532 (P2) |
| CC<sub>33-111</sub> | 29.4 | 8.8 kDa | 25 kDa (2.8) | 1.006 | 0.606 |
| CC<sub>112-185</sub> | 32.9 | 8.9 kDa | 21 kDa (2.4) | 1.036 | 0.526 |
|  |  |  |  | 1.118 (P2) | 1.801 (P2) |
| mLZ<sub>93-171</sub> | 32.2 | 9.1 kDa | 20 kDa (2.2) | 1.072 | 0.586 |
|  |  |  |  | 1.078 (P2) | 1.268 (P2) |

Chi<sup>2</sup> and normalized spatial discrepancy (NSD) values are from *ab initio* modeling. P2 indicates values obtained from *ab initio* modeling when constrained to P2 symmetry.

To the best of our knowledge, the Y-shape topology is unprecedented among both naturally occurring and designed CCs. Modular proteins containing repeat structures are common within the extracellular matrix proteome (Hynes and Naba, 2012), and the addition of the myocilin CC diversifies the modular arrangement. The myocilin CC further expands our appreciation for the versatility of deceptively straightforward CCs and should motivate the design of bio-inspired materials with novel supramolecular arrangements, a highly active research area of protein science (Lupas and Bassler, 2017).

### Identification of parallel tetramer and dimer CC subdomains within Y-shaped molecule

To probe the molecular principles governing the adoption of the Y-shaped structure, we next tested the hypothesis that the Y-shaped molecule apparent from the SEC-SAXS envelope of CC_60-185_ is composed of distinct tetramer and dimer regions corresponding to the two CC sub-regions predicted by Marcoil (Delorenzi and Speed, 2002) and Logicoil (Vincent et al., 2013), residues ~74-102 and ~118-180 (Figure 1). The former stretch escapes detection by other CC prediction programs but even when predicted, assignment of its oligomeric state is of low confidence (not shown). The latter stretch is well predicted as it has regularly spaced Leu residues indicative of the leucine zipper CC subtype.

Biophysical characterization of the Cys-free CC_69-184_ exhibits moderate stability (Table 1), high helical content, and elutes from SEC at a volume consistent with a molecular mass consistent of a tetramer (Supplemental Fig. S3a,b). The addition of Cys_185_ to yield CC_69-185_ increases expression yields and increases thermal stability by 20 °C (Table 1) but secondary and quaternary structures do not change (Supplemental Fig. S3b,c). Non-reducing gel analysis of disulfide bond contributions to the oligomeric state CC_69-185_ reveal only dimers, not higher ordered species (Supplemental Fig. S3d). The SEC-SAXS envelope of CC_69-185_ reveals a P2 symmetric, V-shaped molecule with dimensions 18 nm x 10 nm (Figure 3b, Table 2, Supplemental Fig. S4, Supplemental Fig S5a). The envelope of CC_69-185_ overlays well with the larger CC_60-185_ construct (Supplemental Fig S5b), and appears to be missing the third ‘stem’. We suggest that the stem is not resolved in this construct because, without the disulfide bond from Cys_61_, the protein is susceptible to proteolysis.

To clarify the orientation of CC_69-185,_ relative to the Y-shaped CC_60-185,_ we characterized two additional constructs, CC_33-111_ and CC_112-185_, biochemically and by SEC-SAXS (Supplemental Fig. S6, S7, Table 2). Both proteins are, as expected, helical (Supplemental Fig. S6a,d) but the SEC-SAXS envelopes reveal distinct rods for each constructs (Figure 3c, d). By analysis of its SEC profile, CC_33-111_ is a tetramer (Supplemental Fig. S6b). The corresponding rod is not symmetric, with half of the rod length ~4 nm wide and half ~2 nm wide, perhaps reflecting structural differences in the predicted unstructured N-terminal region and the predicted CC region starting near Gln 75. The SEC profile of CC_112-185_ indicates a dimer (Supplemental Fig. S6e), which is disulfide bonded (Supplemental Fig. S6f,g). CC_112-185_ best matches the length of the arms of the V-shaped CC_69-185_ (Figure 3c, Supplemental Fig. S5c,d). There is a bulge observed approximately in the middle of the CC_112-185_ rod, a feature seen in the pairwise distribution (P(r)) plot of the SEC-SAXS data (Supplemental Fig. S5d, Supplemental Fig. S7h, and see below). These data indicate that CC_60-185_ is composed of two sequentially identical C-terminally disulfide-bonded dimers, which converge at the N-terminal region to form a tetrameric stem (Figure 3e).

### Crystal structure of the 7 C-terminal heptad repeats reveals atypical stabilization for a leucine zipper

Next, we solved a 1.9 Å resolution crystal structure of the C-terminal 7 heptad repeats of the mouse myocilin leucine zipper region, mLZ_122-171_, (residues 136-185 in human myocilin, 80% sequence identity, Figure 4a, Supplemental Table S1). This crystal grew after *in situ* proteolysis of mLZ_93-171,_ a construct nearly identical to CC_112-185_ in sequence and structure (Figure 3c, Supplemental Fig. S8). The susceptibility of mLZ_93-171_ to further proteolysis is not immediately clear, but the presence of a so-called skip residue at Thr89/Thr103 (Figure 4a) suggests the possibility of extended unraveling as seen in myosin (Taylor et al., 2015). The structure of mLZ_122-171,_ which was solved by combining *ab initio* modeling with molecular replacement, reveals the C-terminal disulfide bond (Figure 4b) anticipated from biochemical experiments (Table 1, Figure 2c, Supplemental Fig. S3d, Fig. S6f, g), capping a parallel dimer with a left-handed supercoiled twist (Figure 4c). The two coiled coils in the asymmetric unit are nearly identical to each other, as are the two helices within each coiled coil, with root mean squared deviations (r.m.s.d.) ~1.5 Å (not shown).

**Figure 4.**
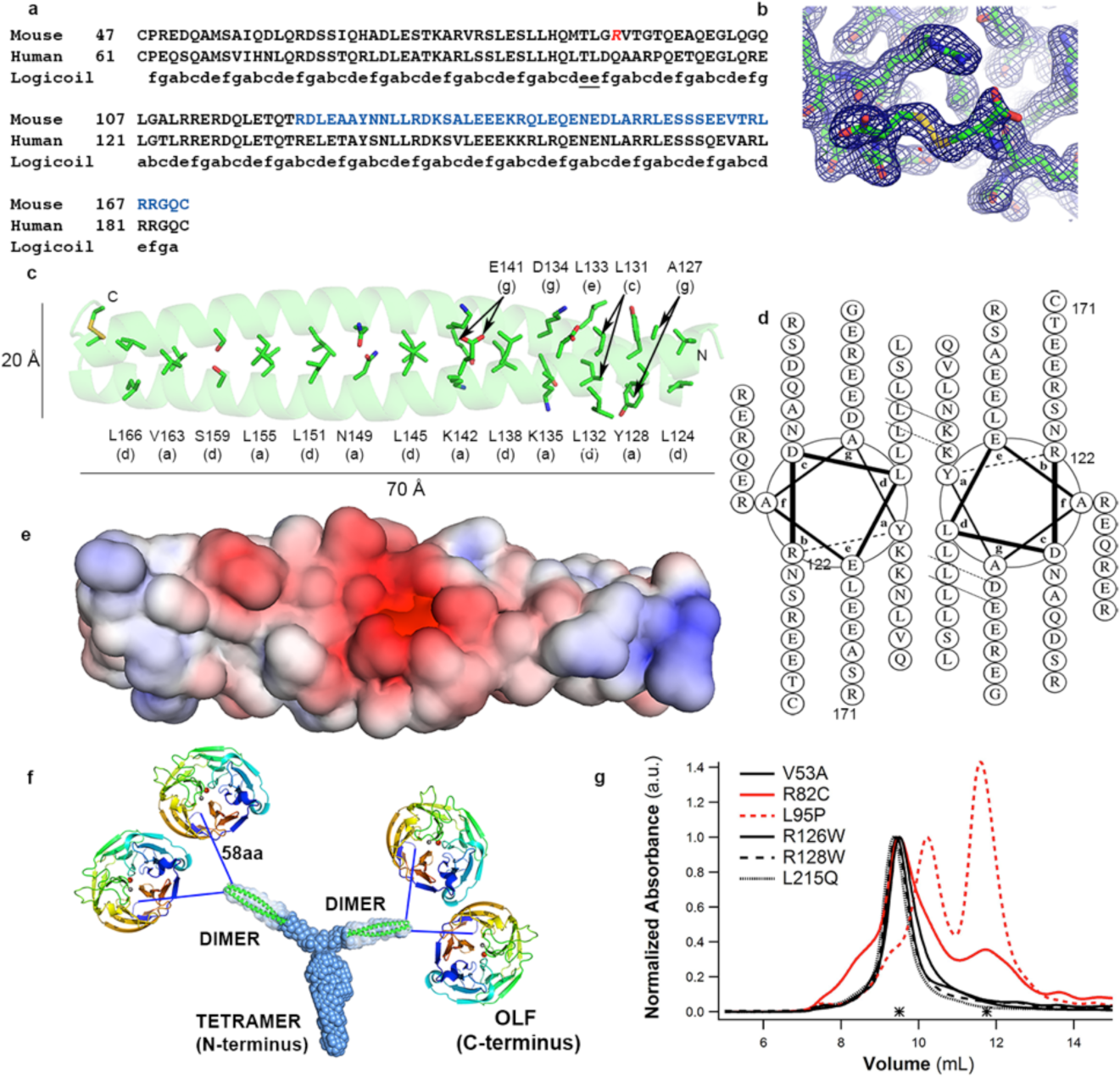
Crystal structure of mLZ_122-171,_ overall myocilin model, and effects of selected glaucoma-relevant variants. (a) Partial sequence alignment of human and mouse myocilin with predicted register of heptad repeats. Red Arg, start of the construct used for crystallization (mLZ_93-171_). Blue stretch, crystallized fragment mLZ_122-171._ (b) Two mLZ_122-171_ helices are covalently tethered by a C-terminal disulfide bond. Final 2Fo-Fc (blue) electron density map is contoured at 1.0 σ and Fo-Fc (red/green) is contoured at ±3.5 σ. (c) Ribbon diagram of mLZ_122-171_ parallel dimer with labeled heptad positions (a) and (d) and disulfide represented as sticks. (d) mLZ_122-171_ in helical wheel representation. (e) Electrostatic surface (negative (red, -5 kT/e-) to positive (blue, + 5 kT/e-)) of mLZ_122-171_ dimer. (f) Overall model of full-length myocilin: Y-shaped dimer-of-dimers composed of an N-terminal parallel tetrameric domain leading to two parallel CC dimers, followed by 58 amino acid long linkers, and C-terminal OLF domains. (g) SEC profile comparison of six NTD_33-227_ glaucoma-relevant variants reveals altered elution and thus quaternary structure for R82C and L95P (red traces).

Analysis of the mLZ_122-171_ structure reveals key deviations from expected general patterns found in the stereotypical “knobs-in-holes” CC (Parry et al., 2008) (Figure 4c, d). Overall, mLZ_122-171_ follows the pattern of a hydrophobic residue, Leu, in position ‘*d*’ except for the penultimate heptad in which it is a polar Ser159. By contrast, fewer than half of the residues in the ‘*a*’ register are hydrophobic. The four remaining residues at ‘*a*’ positions, which are highly conserved among myocilin from human, mouse, rat, fish and cow (Supplementary Figure S9a), are positively charged or polar. Two Lys residues are in positions poised to stabilize negatively charged aspartate and glutamates found one residue N-terminal in the adjacent helix at the ‘*g*’ position, though we note these distances are somewhat long for electrostatic interactions (~3-5 Å). Consensus from prior studies of CCs has suggested that ‘*a*’ positions can tolerate polar residues because they are less buried. Polar residues like Asn are specifically associated with parallel dimers (Mason and Arndt, 2004); in our structure, Asn149 is in position ‘*a*’. Charged residues in ‘*a*’, particularly the frequency of their occurrence is unexpected. One protein design study identified this atypical pairing as important for the specificity of CC interactions among closely related sequences (Grigoryan et al., 2009), although the need for such specificity is not obvious for myocilin. In support of the importance of the *a*-*g* pairing for specificity, and not the traditional ‘*e*’ and ‘*g*’ positions, there are three negatively charged residues in position ‘*e*’ in mLZ_122-171_ and four in ‘*g’*, with just one positively charged residue in ‘*e*’ and ‘*g*’ (Figure 4d). Two of the acidic residues in ‘*g*’, which do not participate in salt bridges with ‘core’ Lys residues in ‘*a*’ positions, are proximal to Asn149 and impart the overall negative electrostatic surface charge of mLZ_122-171_ (Figure 4e). Remarkably, despite these non-canonical repeat features, mLZ_122-171_ overlays well with the closest structural, but distant sequential, homologs, the prototypical leucine zipper GCN4 and geminin (24% and 19% sequence homology across 40 residues, and r.m.s.d.= 1.05 Å and 1.54 Å, respectively, not shown).

### Molecular dynamics (MD) simulations suggest the potential for mLZ_122-171_ to adopt an alternative structure

After 50 ns of equilibration at 310 K, the helical structures of mLZ_122-171_ and the disulfide-deletion variant mLZ_122-170_ remain essentially unchanged (Supplemental Fig. S10a). However, after 100 ns at 400 K, a bend emerges in the structure of mLZ_122-171_ near the C-terminus (Ser159-Gln170), concomitant with a shift in paired interactions between oppositely charged residues on either helix (Supplemental Fig. S10b,c). This region of the leucine zipper includes paired Ser159 residues located in the heptad ‘*d*’ position as well as other polar/charged residues. Without the disulfide and consistent with stability measurements (Table 1), the mLZ_122-170_ dimer is considerably more susceptible to unfolding. For example, in the 400 K simulation, the distinct kinked structure was not formed. In steered molecular dynamics simulations at 310 K in which the centers-of-mass of each coil were pulled apart at a constant velocity of 1 Å/ns for 10 ns, the C-terminal ends separated completely (Supplemental Fig. S10d). In line with its more hydrophobic character, the N-terminal region of mLZ_122-171_ and mLZ_122-170_ dimers remained unaltered in all simulations. In sum, the disulfide bond at the C-terminus of the myocilin leucine zipper appears to limit the extent to which each protomer can unfold and the prevalence of complementary charged residues allows the coil to adopt an alternative structure. Such resistance to unfolding is likely a feature that helps myocilin cope with the regular mechanical distortions of the TM ocular tissue (Acott et al., 2014).

### Model of full-length myocilin

With the molecular structures of the myocilin CC and OLF domains (Donegan et al., 2015) in hand, we propose the following model of full-length myocilin (Figure 4f). Myocilin adopts a distinct oligomeric state defined by the dimer-stabilizing trio of Cys residues 47, 61, and 185, and the CCs, which confer a unique Y-shaped architecture of a branched dimer-of-dimers tetramer. We found no evidence for higher ordered states for myocilin seen in previous studies (Fautsch et al., 2004; Russell et al., 2001). Higher ordered states might form *in vivo*, perhaps mediated by a yet-unidentified binding partner not present in our purified samples. It is also possible that high molecular weight myocilin was observed because myocilin in prior studies was misfolded, either due to incomplete posttranslational processing of the signal sequence or due to handling of antioxidant aqueous humor fluid (Richer and Rose, 1998) *ex vivo*, both of which could lead to aberrant inter-molecular disulfide bond formation. Two parallel dimers emerge at an obtuse angle from the N-terminal tetrameric stalk, imparting multivalency to the otherwise monomeric C-terminal OLF domain, likely to increase avidity and space OLF pairs apart from one another. Based on our constructs the transition from tetramer to the two dimers begins between Ser69 and Glu112, and most likely includes the lower-CC probability region containing the skip residue, Thr103 (Figure 4a, Supplemental Figure S9a). Aside from structural regions, myocilin also contains lengthy, linker regions on both N- and C-terminal ends of the structural Y, the functional/structural significance of which is not currently known.

### Implications for other olfactomedin family members

The high level of sequence conservation across myocilin-like subfamily 3 members (Supplemental Fig. S9b) indicates the Y-shape architecture is shared across eukaryotes. The topology of myocilins is distinct from the dimer-of-dimers V of olfactomedin-1 (subfamily 1), however, likely a result of the different coil prediction pattern (Figure 1) and low overall sequence similarity. For example, angle of the V of olfactomedin-1 appears acute, not obtuse like measured for CC_69-185._ In addition, the C-terminal region of the olfactomedin-1 CC, which harbors a disulfide-forming Cys found within the CC (Pronker et al., 2015), is not aligned with the CC-capping residue Cys 185 in myocilin (Figure 4b, Supplemental Fig. S1b). In olfactomedin-1, the end of the disulfide-harboring CC is immediately adjacent to the start of the OLF domain (Pronker et al., 2015) whereas in myocilin Cys185 is 60 residues away from the start of the structural OLF at Cys245. Whether the remaining three CC-containing olfactomedin subfamilies adopt a dimer-of-dimers and to what extent they are similar to myocilin or olfactomedin-1 remains to be seen. Other open questions across the olfactomedin family surround the functional consequences of the unusual olfactomedin topologies. A naive interpretation would suggest the CCs are tailored “molecular rulers,” to space pairs of OLF domains, but mechanotransduction or as molecular scaffolds for complex protein-protein interactions are intriguing alternative possibilities.

### Structural defects imparted by single nucleotide polymorphism and glaucoma-associated myocilin variants

In contrast to mutations found within the myocilin OLF domain, many of which are associated with early-onset and familial glaucoma (Hewitt et al., 2006), the vast majority of mutations in the N-terminal module have been identified via case-control genome sequencing studies. As such, their annotation as pathogenic is predicted statistically. The extent to which the toxic gain-of-function hypothesis for myocilin-associated glaucoma (misfolding/non-secretion/aggregation) holds for mutations found beyond OLF is unclear because such variants have not been systematically studied. Just one study investigated cellular trafficking and adhesion properties of two annotated N-terminal disease variants (R82C and R126W, a rare late-onset familial mutation (Faucher et al., 2002)) and one neutral (L95P) mutant (Gobeil et al., 2006). Compared to most OLF-resident mutations, which were retained intracellularly (Liu and Vollrath, 2004; Vollrath and Liu, 2006; Zhou and Vollrath, 1999), all three N-terminal variants tested (Gobeil et al., 2006) were secreted, but for R82C and L95P variants, partial intracellular sequestration was observed in a cell-based assay. R82C- and L95P-mutant myocilins also lost adhesion properties in a cell assay (Gobeil et al., 2006), but R126W did not.

On the basis of the examined wild-type CC constructs, which exhibit reversible thermal unfolding (Figure 2a, Supplementary Fig. S3a), it is apparent that the CC domain is not intrinsically prone to aggregation like OLF, but disease-associated N-terminal mutants might still misfold by exhibiting structural anomalies without aggregation. To test our hypothesis, we biophysically characterized six NTD_33-227_ variants (Table 1, Figure 4g, Supplemental Fig. S11). Linker-resident, conservative substitutions V53A and L215Q are annotated as glaucomatous (Hewitt et al., 2007). Biophysically, however, they are indistinguishable from wild-type in stability, thermal unfolding reversibility, and SEC elution profile, indicating no major structural changes have occurred. In the dimer region of the CC, the two changes R126W and R128W, the latter annotated as neutral, are expected to be poorly tolerated. This is especially true for R128W, in which the Arg appears to fall in the electrostatically positive ‘*a*’ position for pairing with a negatively charged Glu ‘*g*’ in an adjacent helix. Surprisingly, both variants behaved as wild-type. In the tetramer region, which has thus far eluded crystallization, similarly non-conservative substitutions R82C and L95P previously mentioned, were also not expected to be well tolerated. Both of these variants exhibited wild-type like stability and reversible thermal unfolding, with no evidence for aggregation (Supplemental Fig. S11). The elution profiles from SEC differ from wild-type, however, in that they are both mixtures of wild-type and smaller, non-native species (Figure 4g).

In sum, our characterization of myocilin reveals a new topology for CCs, an unprecedented branched Y-shaped protein structure. The tetramer region is sensitive to mutation, as changes in quaternary structure were observed for R82C and L95P, with a consequence of non-native cellular secretion and poor cellular adhesion observed previously. Further studies will be required to assess whether both variants are glaucoma-causing, and to what extent gain-of-function (misfolding, intracellular sequestration) and/or loss-of-function (adhesion, mechanical properties, other) mechanisms are at play. By contrast, the dimer region appears relatively resilient to mutation and unfolding, and may intentionally access multiple distinct conformations. Documented linker-resident mutations, which are not expected to affect the quaternary structure nor located at positions with high likelihood for posttranslational modification, may not be causative for glaucoma. Beyond expanding the scope of naturally occurring protein structures, defects associated with new variants identified in the population are synergistic with efforts to identify at-risk patients to manage glaucoma prior to onset of vision loss (Souzeau et al., 2016) and to develop personalized medicine for glaucoma.

## Materials and Methods

### Molecular Cloning, Protein Expression, and Purification

Constructs of N-terminal portions of human myocilin were obtained by subcloning from an *E. coli* codon-optimized construct of full-length human myocilin (DNA 2.0) as described in Supplemental Methods. Proteins were expressed in *E. coli* BL21(DE3)plysS cells using the same protocol. A single transformed colony was inoculated into a selective 150-250 mL Luria-Bertani culture (LB, Fisher, supplemented with 50 μg/mL kanamycin and 34 μg/mL chloramphenicol) and agitated at 225 r.p.m., 37 °C, overnight (16-24 hours). A starter culture (20 mL) was used to inoculate 1L of Superior Broth media (U.S. Biological), which was then agitated at 225 r.p.m., 37 °C until an optical density at 600 nm of 1.0-2.0 was reached. Protein expression was induced with 0.5 mM ispropyl β-D-thiogalactopyranoside (IPTG) at 18 °C and induced cells were grown overnight (16-20 hours) by agitation at 225 r.p.m., 18 °C. Cells were harvested by centrifugation and flash frozen in liquid nitrogen for storage at -80 °C. Cells were lysed by French Press, protein constructs purified by Ni^2+^-affinity and size exclusion chromatography, and further evaluated as detailed in Supplemental Methods.

### Circular Dichroism

Far-UV circular dichroism (CD) spectra and thermal melts were acquired on a Jasco J-815 spectropolarimeter equipped with a Jasco PTC-4245/15 temperature control system. Protein samples at a concentration range of 5-30 μM were measured in 50 mM Tris pH 7.5, 200 mM NaCl buffer at 4 °C. Scans were measured from 200 nm to 300 nm at a rate of 200 nm/min and a data pitch of 1 nm using a 0.1 cm cuvette. Each measurement was blank-subtracted with buffer and is an average of 10 scans. Far-UV thermal melts were performed from 4 to 80 °C at a rate of 1 °C min^−1^ increase in temperature and a data pitch of 2 °C. Ten scans from 300 to 200 nm at a 200-nm min^−1^ scan rate were averaged for each temperature. The T_m_ was determined by plotting the ratio of values recorded at 208 nm and 222 nm versus temperature and analyzed by Boltzmann Sigmoid analysis using Igor Pro.

### SEC-SAXS

SEC–SAXS experiments were performed at BioCAT (beamline 18ID, Advanced Photon Source at Argonne National Labs); details of the experimental set up and data analysis are presented in Supplemental Methods.

### Crystallization, data collection, structure determination

Crystals of mouse LZ_93-171_ (later determined to be mouse LZ_122-171)_ at 20 mg/mL in 10 mM Hepes pH 7.5 No NaCl grew after 90 days in 0.1 M ammonium acetate, 0.1 M BisTris pH 5.5, and 17% PEG 10,000 from a Hampton Index, HR2-134, sparse matrix sitting drop tray. Crystals were cryoprotected in 0.1 M ammonium acetate, 0.1 M BisTris pH 5.5, 25% PEG 10,000, and 25% glycerol. Data were collected at the Southeast Regional Collaborative Access Team (SER-CAT) 22-ID beamline and processed using HKL-2000 (Otwinowski and Minor, 1997). An initial molecular replacement solution was found using AMPLE (Bibby et al., 2012), which consisted of three ~3 kDa helical poly-alanine chains with reasonable electron density. Using one of these ~3 kDa chains another round of molecular replacement was performed with Phaser (McCoy et al., 2007) but with a search criteria of eight molecules in the asymmetric unit as suggested by analysis of Matthews Coefficient (Kantardjieff and Rupp, 2003). Visualization of neighboring unit cells in Coot (Emsley et al., 2010) suggested a probable connection of pairs of chains, leading to the final solution of four ~6 kDa poly-alanine chains in the asymmetric unit. The poly-alanine chains were iteratively built and refined using Coot (Emsley et al., 2010) and Phenix.refine (Afonine et al., 2012) by slowly replacing alanines with respective amino acids to fit the electron density. The final model consists of mouse residues 122-171. The structure has been deposited to the PDB with accession number 5VR2.

### Molecular dynamics (MD) simulations

Two structures were prepared for MD simulations: the dimer of mLZ_122-171_ both with (WT) and mLZ_122-171_ without (ΔC171) the C-terminal cysteines. Each structure was solvated in a water box of dimensions 105 × 105 × 105 Å^3^ using VMD (Humphrey et al., 1996). The system was neutralized with 0.15 M NaCl, resulting in a total of ~110,000 atoms for each system. Molecular dynamics simulations were carried out using both the NAMD 2.12 (Phillips et al., 2005) and Amber16 (Case et al., 2016) programs with the CHARMM36 all-atom force field (Best et al., 2012a; Best et al., 2012b). The temperature was fixed using Langevin dynamics; the pressure was kept constant at 1 atm using the Langevin piston method (Feller et al., 1995). The equations of motion were integrated using the RESPA multiple time-step algorithm with a time step of 2 fs used for all bonded interactions, 2 fs for short-range non-bonded interactions, and 4 fs for long-range electrostatic interactions. Long-range electrostatic interactions were calculated using the particle-mesh Ewald method (Darden et al., 1993). Bonds involving hydrogen atoms were constrained to their equilibrium length employing the Rattle algorithm (Andersen, 1983).

### Bioinformatics

Bioinformatics programs used to analyze sequences and structures are listed in Supplemental Methods.

## Acknowledgements

This work was funded by NIH R01EY021205 to R.L.L. Use of the Advanced Photon Source was supported by the U. S. Department of Energy, Office of Science, Office of Basic Energy Sciences, under Contract No. W-31-109-Eng-38. Work performed at Bio-CAT was supported by NIH NIGMS 9P41 GM103622. Use of the Pilatus 3 1M detector was provided by NIGMS 1S10OD018090-01. We thank Srinivas Chakravarthy for SEC-SAXS data collection.

## Author Contributions

S.E.H., R.L.L., J.C.G. designed experiments, interpreted results, wrote, and edited the manuscript. S.E.H., E.N., R.K.D., A.H. conducted experiments.

## Supplemental Materials and Methods

### Molecular Cloning and Protein Expression

Constructs of N-terminal portions of human myocilin were obtained by subcloning from an *E. coli* codon-optimized construct of full-length human myocilin (DNA 2.0). Primers for NTD_33-227_, CC_60-185_, CC_69-184_, CC_33-111_, and CC_112-184_, were designed to incorporate a C-terminal TAA stop codon as listed in Supplemental Table S2. Amplified DNA was purified via a Promega PCR purification procedure, treated with T4 DNA polymerase, and annealed into the pET-30 Xa/LIC vector following the manufacturer’s directions. Cys-to-Ser variants of NTD_33-227_ (C_61_S/C_185_S, C_47_S/C_185_S, C_47_S/C_61_S, C_47_S/C_61_S/C_185_S), CC_69-185_, CC_112-185_, and NTD_33-227_ disease-causing variants were generated by site directed mutagenesis (QuikChange Stratagene mutagenesis kit, primers listed in Supplemental Table S2). Plasmids were propagated by *E. coli* NovaBlue GigaSingles and the correct in-frame DNA sequence was confirmed (MWG Operon). The plasmid for Mouse LZ_55-171,_ designed to be identical to the pET-30 Xa/LIC vector with a Factor Xa protease cleavage site was purchased from Genscript. After proteins were expressed in *E. coli* BL21(DE3)plysS cells, they were purified as detailed below.

**Protein Purification**

#### NTD_33-227_ (WT)

Cell pellets (20 g) were resuspended using a serological pipette into 40 mL Ni^2+^-affinity purification wash buffer (50 mM Hepes pH 7.5, 200 mM NaCl, 40 mM imidazole, 10% glycerol) supplemented with 2 EDTA-free Roche Protease Inhibitor Cocktail tablets. Cells were lysed by 2-3 passages through a French press (12,000 psi). To increase protein yield, 1% Trion X-100 was added and lysed cells were rocked gently in a cold room for 1 hour. Cellular debris was removed by ultracentrifugation at 130,000 × g, 4 °C for 1 hour and the supernatant loaded onto a 1 mL Ni^2+^-affinity purification column (GE Healthcare) using an AKTA purification system (GE Healthcare). After 120 column-volumes of wash buffer was loaded to remove contaminating proteins, NTD_33-227_ was eluted with a gradient from 40-500mM imidazole using a mixture of wash buffer and elution buffer composed of 50mM Hepes pH 7.5, 200mM NaCl, 500 mM imidazole, 10% glycerol. Fractions containing _NTD33-227_ were concentrated using Amicon 100K MWCO filters (EMD Millipore) and loaded onto a Superdex 200 10/300 GL (GE Healthcare) size exclusion column (SEC) equilibrated with gel filtration buffer (50 mM Hepes pH 7.5, 200 mM NaCl, 10% glycerol). The fractions containing NTD_33-227_ were identified by SDS-PAGE analysis (15% polyacrylamide) with Coomassie staining and found to have a contaminating band of DnaK, a result confirmed by trypsin digest mass spectrometry (Supplemental Fig. S12a, Systems Mass Spectrometry Core Facility, Georgia Tech). Taking advantage of the finding that NTD_33-227_ can unfold reversibly (Figure 2a), an on-column unfolding refolding protocol (see below) was used to remove the contaminating DnaK for selected preparations (e.g. Figure 2 and NTD_33-227_ Cys-to-Ser variants below). Removal of DnaK did not alter the elution profile of NTD_33-227_ from Superdex 200 (Supplemental Fig. S12b,c). Protein concentration was estimated by absorbance at 280 nm using a molecular extinction coefficient (8,605 M^−1^cm^−1^) and a molecular weight (27,014.8 g/mol) calculated by ExPASy ProtParam (Gasteiger et al., 2005)

#### NTD_33-227_ Cys-to-Ser Variants (C47S/C61S, C61S/C185S, C47S/C185S, C47S/C61S/C185S)

Cell pellets (20 g) were lysed using the Triton X-100 step and purified by Ni^2+^-affinity column chromatography as outlined above. Fractions containing NTD_33-227_ were concentrated to 1 mL using Amicon 100K MWCO filters (EMD Millipore) followed by dilution into 50mL urea unfolding buffer (50 mM Hepes pH 7.5, 500 mM NaCl, 8 M Urea). Unfolded NTD_33-227_ was reloaded onto the Ni^2+^-affinity column, washed with 20 column volumes of urea unfolding buffer to remove contaminating DnaK, and then refolded on the column by a gradient into refolding buffer (50mM Hepes pH 7.5, 200mM NaCl, 10% glycerol). Folded NTD_33-227_ was eluted by a gradient from 0-500 mM imidazole also containing 50mM Hepes pH 7.5, 200mM NaCl, 10% glycerol, fractions containing NTD_33-227_ were concentrated using Amicon 30K MWCO filters (EMD Millipore), and samples were then purified by Superdex 200 10/300 GL SEC as described above. Purified proteins were assessed and quantified as for WT NTD_33-227_.

#### Annotated NTD_33-227_ Variants (V53A, R82C, L95P, R126W, R128W, L215Q)

Cell pellets (10 g) were resuspended using a serological pipette into 20 mL Ni^2+^-affinity purification wash buffer supplemented with 1 EDTA-free Roche Protease Inhibitor Cocktail tablet. Cells were lysed, debris removed, and protein purified as described for NTD33-227. Fractions containing NTD_33-227_ were assessed by SDS-PAGE analysis, concentrated with a 100K MWCO filter, and re-loaded onto a Superdex 200 10/300 GL equilibrated with gel filtration buffer to obtain the final presented chromatogram.

#### CC_60-185_

Cell pellets (20 g) were resuspended, lysed, and purified by Ni^2+^-affinity and SEC as for NTD_33-227_, except that the initial cell resuspension buffer was further supplemented with 1 mL of 10 mg/mL DNase, 200 μL of 1 M CaCl2, 200 *μ*L of 1 M MgCl_2_ and the Triton X-100 step was not used. To remove tags, purified CC_60-185_ was incubated with Factor Xa (Roche, ~50:1 mass ratio) plus ~5-10 mM CaCl_2_ for 96 hours at 4 °C. Digested CC_60-185_ was diluted ~20 fold into heparin wash buffer (20 mM Hepes, pH 7.5, 10% glycerol), loaded onto a 5 mL HiTrap Heparin HP column (GE Healthcare), and eluted by a gradient from 0-1 M NaCl with buffers also containing 20 mM Hepes pH 7.5, 10% glycerol, which also removed Factor Xa and DnaK contaminants. Heparin elution fractions were assessed by SDS-PAGE analysis as above, and those containing CC_60-185_ were concentrated with Amicon 10K MWCO filters and loaded onto a HiLoad 16/600 Superdex 75 prep grade column (GE Healthcare) equilibrated with gel filtration buffer. Fractions corresponding to a molecular weight of ~60 kDa based on a standard calibration curve of the SEC column, containing pure CC_60-185_ as assessed by SDS-PAGE analysis with Coomassie staining, were concentrated in an Amicon 10K MWCO filter. Protein concentration was measured by absorbance at 280 nm using a molecular extinction coefficient (1,490 M^−1^cm^−1^) and a molecular weight (14,643 g/mol) calculated by Expasy ProtParam.

#### CC_69-185_

Cell pellets (20 g) were resuspended and lysed as for CC_60-185_ except that the 1% Triton X-100 procedure was performed as for WT NTD_33-227._ Purification by Ni^2+^ affinity and SEC proceeded as described for WT NTD_33-227._ After assessment by SDS-PAGE analysis, fractions containing CC_69-185_ were digested and purified by heparin affinity and Superdex 75 following procedures for CC_60-185_ Fractions corresponding to a molecular weight of ~50 kDa based on an SEC standard calibration curve containing pure CC_69-185_ as assessed by SDS-PAGE analysis with Coomassie staining were concentrated in an Amicon 10K MWCO filter. Protein concentration was measured by absorbance at 280 nm using a molecular extinction coefficient (1,490 M^−1^cm^−1)^ and a molecular weight (13,681 g/mol) calculated by Expasy ProtParam.

#### CC_33-111_

Cell pellets (20 g) were resuspended, lysed, and purified as described for CC_60-185_ except Amicon 10K MWCO filters were used to concentrate the protein. Following assessment by SDSPAGE analysis, fractions containing pure CC_33-111_ were incubated with Factor Xa (Roche, ~50:1 mass ratio) plus ~5-10 mM CaCl_2_ for 96 hours at 4 °C. Digested CC_33-111_ was concentrated with two Amicon 10K MWCO filters and loaded onto a HiLoad 16/600 Superdex 75 prep grade column (GE Healthcare) equilibrated with gel filtration buffer. Fractions corresponding to a molecular weight of 40 kDa based on a standard calibration curve containing pure CC_33-111_ as assessed by SDS-PAGE analysis with Coomassie staining were concentrated in an Amicon 10K MWCO filter. Protein concentration was measured by absorbance at 280 nm using a molecular extinction coefficient (1,490 M^−1^cm^−1^) and a molecular weight (8,801 g/mol) calculated by ExPasy ProtParam.

#### CC_112-185_

Cell pellets (20 g) were resuspended, lysed, and purified as described for CC_33-111_. Purified CC_112-185_ was digested and further purified using the procedure described for CC_60-185_. Fractions corresponding to a molecular weight of 23 kDa based on a calibration curve for SEC containing pure CC_112-185_ as assessed by SDS-PAGE analysis were concentrated in an Amicon 10K MWCO filter. Protein concentration was measured by absorbance at 280 nm using a molecular extinction coefficient (1,490 M^−1^cm^−1^) and a molecular weight (8,858 g/mol) calculated by Expasy ProtParam.

#### Mouse LZ_55-171_ and LZ_93-171_

Cell pellets (20 g) were resuspended, lysed, and purified as described for CC_60-85_ except 100K MWCO filtration devices were used to concentrate the protein. After assessment by SDSPAGE, fractions containing pure mouse LZ_55-171_ were incubated with Factor Xa (Roche, ~50:1 mass ratio) plus ~5-10 mM CaCl_2_ for 72 hours at 4 °C. Digested mouse LZ_55-171_ was diluted ~30 fold into heparin wash buffer (20 mM Hepes, pH 7.5, 10% glycerol), loaded onto a 5 mL HiTrap Heparin HP column (GE Healthcare). Unlike the human constructs, mouse LZ_55-171_ did not bind the heparin column, so flow through fractions were concentrated with Amicon 10K MWCO filters and loaded onto a HiLoad 16/600 Superdex 75 prep grade column (GE Healthcare) equilibrated with gel filtration buffer. The majority of protein was found in fractions corresponding to a molecular weight of ~30 kDa based on a calibration curve; however, after assessment by SDS-PAGE the dominant band on the gel was below the predicted 13,500 g/mol molecular weight for cleaved mouse LZ_55-171._ Mass spectrometry (Supplemental Fig. S12d) confirmed mouse LZ_55-171_ was cleaved by Factor Xa after a second Gly-Arg amino acid amino acid within the amino acid sequence, yielding mouse LZ_93-171_ used in subsequent experiments. Fractions containing mouse LZ_93-171_ were concentrated in an Amicon 10K MWCO filter. Protein concentration was measured by absorbance at 280 nm using a molecular extinction coefficient (1,490 M^−1^cm^−1^) and a molecular weight (9,144 g/mol) calculated by ExPASy ProtParam for mouse LZ_93-117_.

### SEC-SAXS

SEC–SAXS experiments were performed at BioCAT (beamline 18ID, Advanced Photon Source at Argonne National Labs). The camera included a focused 12 KeV (1.03 Å) X-ray beam, a 1.5 mm quartz capillary sample cell, a sample to detector distance of ~3.5 m, and a Pilatus 3 1M detector (Dectris). The Q-range sampled was ~0.0042–0.4 Å–1. A size-exclusion chromatography setup in line with the SAXS camera was used to separate the sample from potential aggregates, breakdown products and other contaminants using a Superdex 75 10/300 GL (GE Healthcare Lifesciences) column. The elution trajectory after the UV monitor was redirected to the SAXS sample flow-cell. Exposures (1 s) were collected every 2 s during the gel-filtration chromatography run. Data reduction to generate the I(q) vs. q curves for all constructs except CC_60-185_ was performed with the BioCAT beam line specific pipeline, which uses the ATSAS program suite(Petoukhov et al., 2012). For CC_60-185_ multiple I(q) vs. q curves were manually generated and inspected to optimize frame choice, eliminating frames for which signal was being derived from aggregated protein. Exposures flanking the elution peak were averaged and used as the buffer curve for each run, and the exposures corresponding to the UV peak on the chromatogram were treated as protein plus buffer curves. Data were corrected for background scattering by subtracting the buffer curve from protein plus buffer curves. The radius of gyration (R_g_) was calculated using Guinier approximation and a P(r) curve, both of which were done using PRIMUS (Konarev et al., 2003). D_max_ was also calculated using the P(r) curve and the output was used to calculate molecular envelopes using DAMMIN (Svergun, 1999) The envelopes were then averaged using DAMAVER and DAMFILT (Volkov and Svergun, 2003). For CC_69-185_, CC_112-185_, and mLZ_93-171_ results of imposing two-fold symmetry (P2) was compared to results with no imposed symmetry (P1). For all constructs and symmetry constraints presented, at least 9 out of the 10 envelopes were accepted by DAMAVER on the basis of the basis of the Normalized Spatial Discrepancy (NSD). SUPCOMB(Kozin and Svergun, 2001) was used to overlay envelopes of differing symmetry, compare CC_60-185_ with CC_69-185,_ and compare CC_112-185_ compared to mLZ_93-171_ (Supplemental Figure S5 and Supplemental Figure S8e,f). Overlay of CC_112-185_ and CC_33-111_ (and Figure 3e) was performed manually within the PyMOL Molecular Graphic System v. 1.7.6.0. Final *ab initio* models were rendered using Visual Molecular Dynamics (Humphrey et al., 1996). Molecular masses were calculated using SAXS MolW2 (Fischer et al., 2010).

### Bioinformatics

Multiple sequence alignments were generated in CLUSTAL Omega (Sievers and Higgins, 2014). Myocilin sequences from multiple organisms were identified using the human myocilin OLF domain sequence as a query within BLAST (Altschul et al., 1990). The list was manually culled to remove partial sequences and non-myocilin proteins, which were identified either by annotation or by non-myocilin sequence stretches recognized by us through other related structures of OLF domain proteins (Hill et al., 2015). Then the OLF domains were deleted beginning with the first structural residue, Cys (Donegan et al., 2015) leaving 75 N-terminal sequences intact. After multiple sequence alignments, the consensus sequence was visualized using Skylign (Wheeler et al., 2014). Coiled coil prediction servers Marcoil (Delorenzi and Speed, 2002) and Logicoil (Vincent et al., 2013) were utilized to generate models for the coiled-coil domain of myocilin; a consensus cutoff of 80% probability was used to label the limits of the domains presented in Figure 1. Logicoil was used to obtain the coil register for Figure 4a and DrawCoil 1.0 (Grigoryan and Keating, 2008) was used to generate the image presented in Figure 4d. A structure similarity search was conducted with PDBeFOLD (Krissinel and Henrick, 2005) and sequence similarity searches limited to CCs were conducted using CC+ Builder (Testa et al., 2009). Structures were aligned using the secondary structure matching algorithm (Krissinel and Henrick, 2004) to obtain r.m.s.d. values in Coot. Electrostatic surfaces were generated using default options within the PDB2PQR web server (Dolinsky et al., 2007) and the APBS plugin (Baker et al., 2001) within PyMOL.

## Supplemental Figures

**Supplemental Figure S1.**
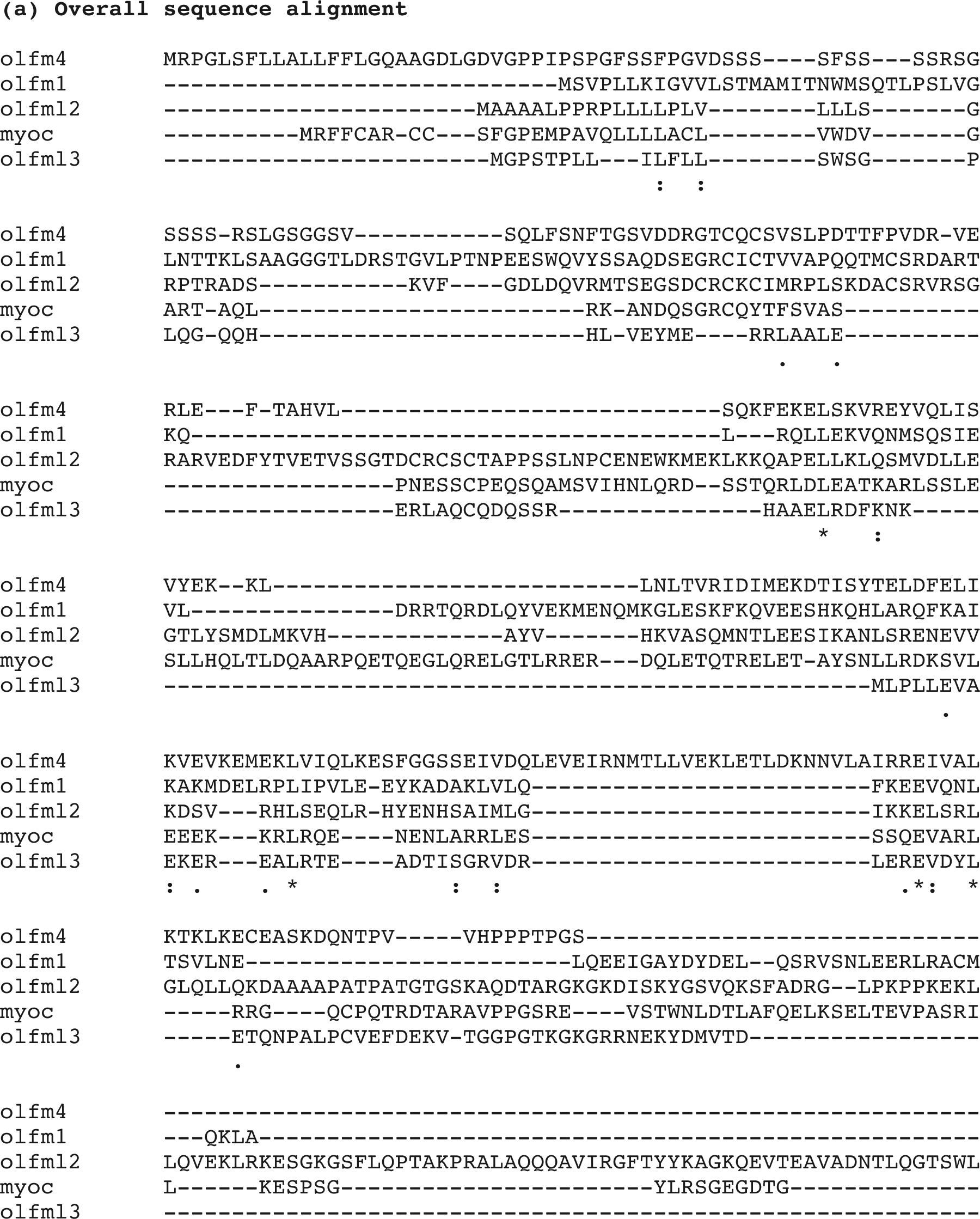
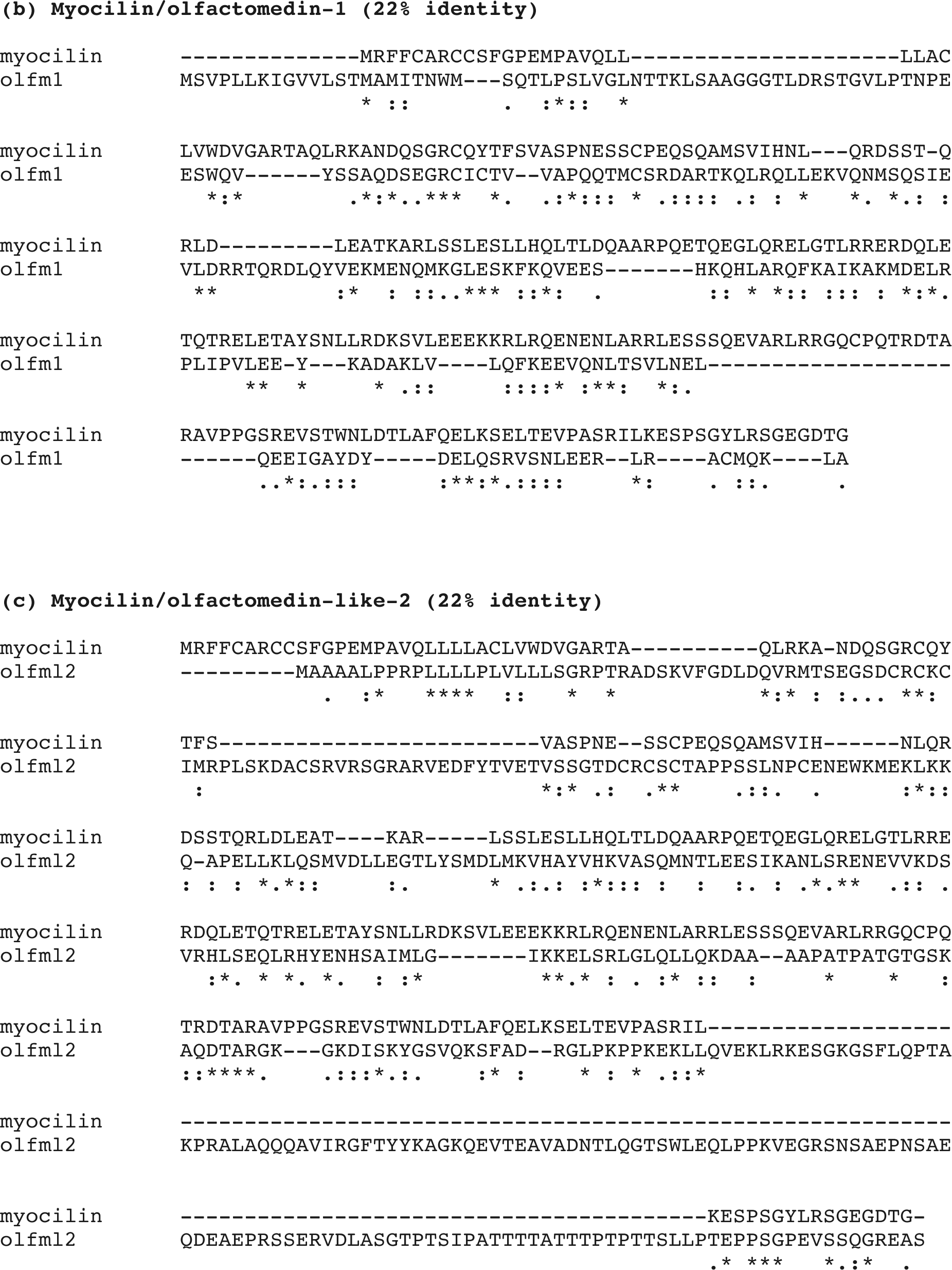
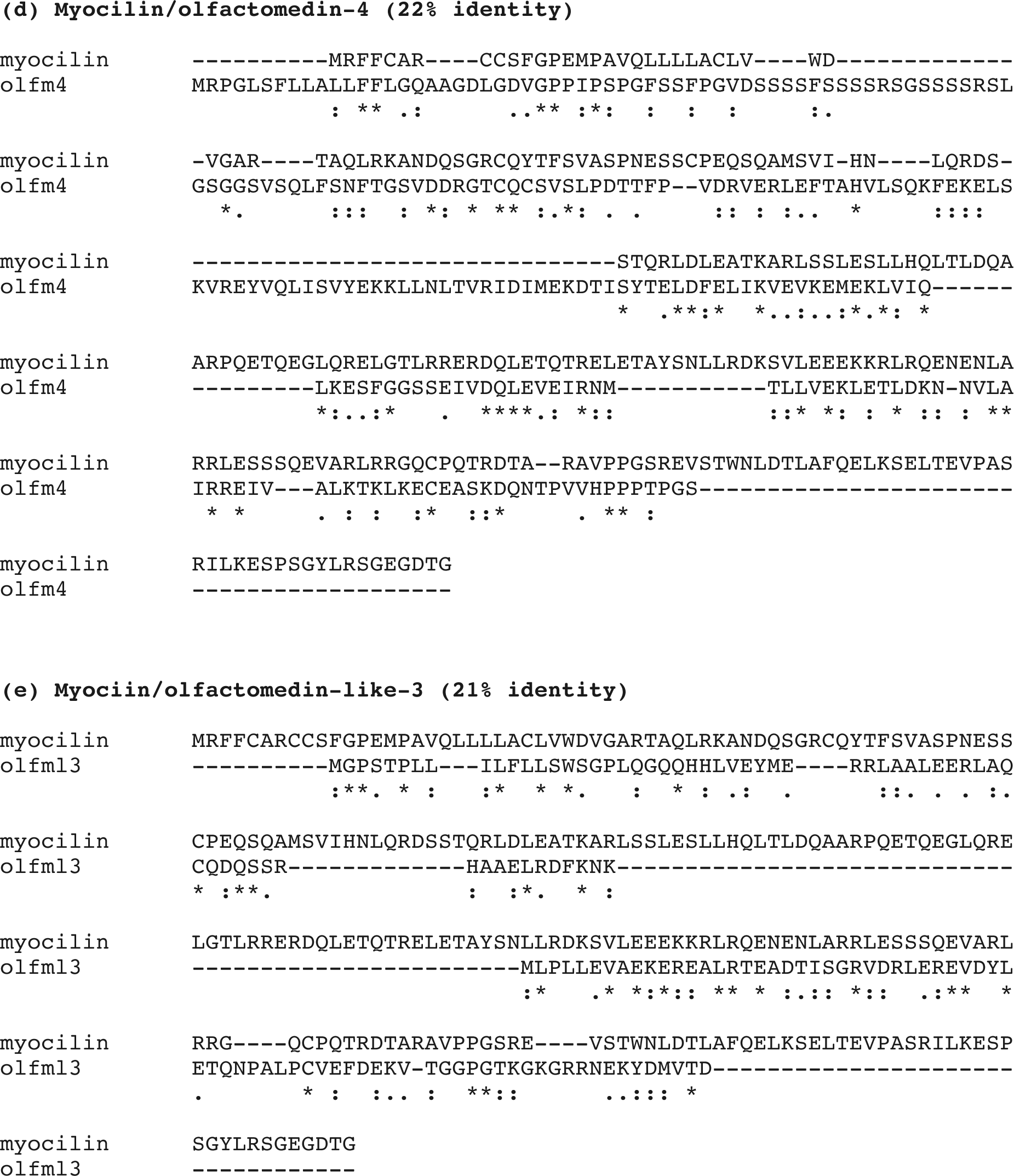
Sequence alignments of CC-containing regions of olfactomedin-containing proteins. Related to Fig. 1. (a) Overall sequence alignment of CC-containing regions of human proteins found within different olfactomedin subfamilies. (b) Pairwise sequence alignment of myocilin and olfactomedin-1. (c) Pairwise sequence alignment of myocilin and olfactomedin-like-2. (d) Pairwise sequence alignment of myocilin and olfactomedin-4. (e) Pairwise sequence alignment of myocilin and olfactomedin-like-3.

**Supplemental Figure S2.**
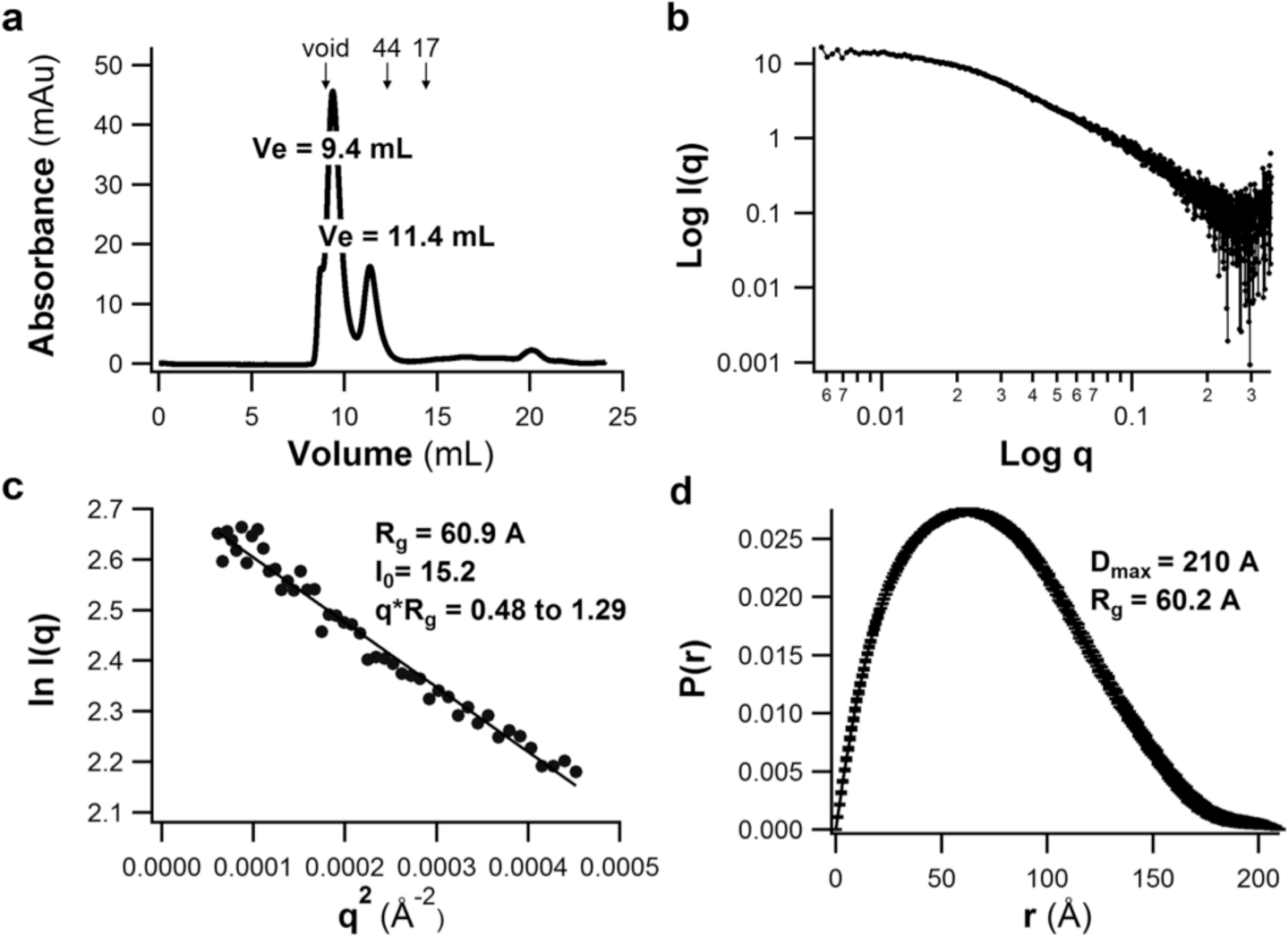
SEC-SAXS analysis of CC_60-185._ Related to Fig. 3. (a) Superdex-75 GL trace of CC_60-185_ obtained at the beamline reveals two species, one in the void volume and another with a molecular weight of 70 kDa based on a standard calibration curve. Given the calculated mass of 14.6 kDa/monomer, the second species is a 4 or 5-mer. Further SAXS analysis focused exclusively on this second species. (b) Scattering intensity profile. I(q): scattered intensity, q:scattering vector. (c) Guinier plot with calculated radius of gyration (Rg). (d) Pair wise distribution plot with calculated maximum particle size (Dmax) and radius of gyration (Rg).

**Supplemental Figure S3.**
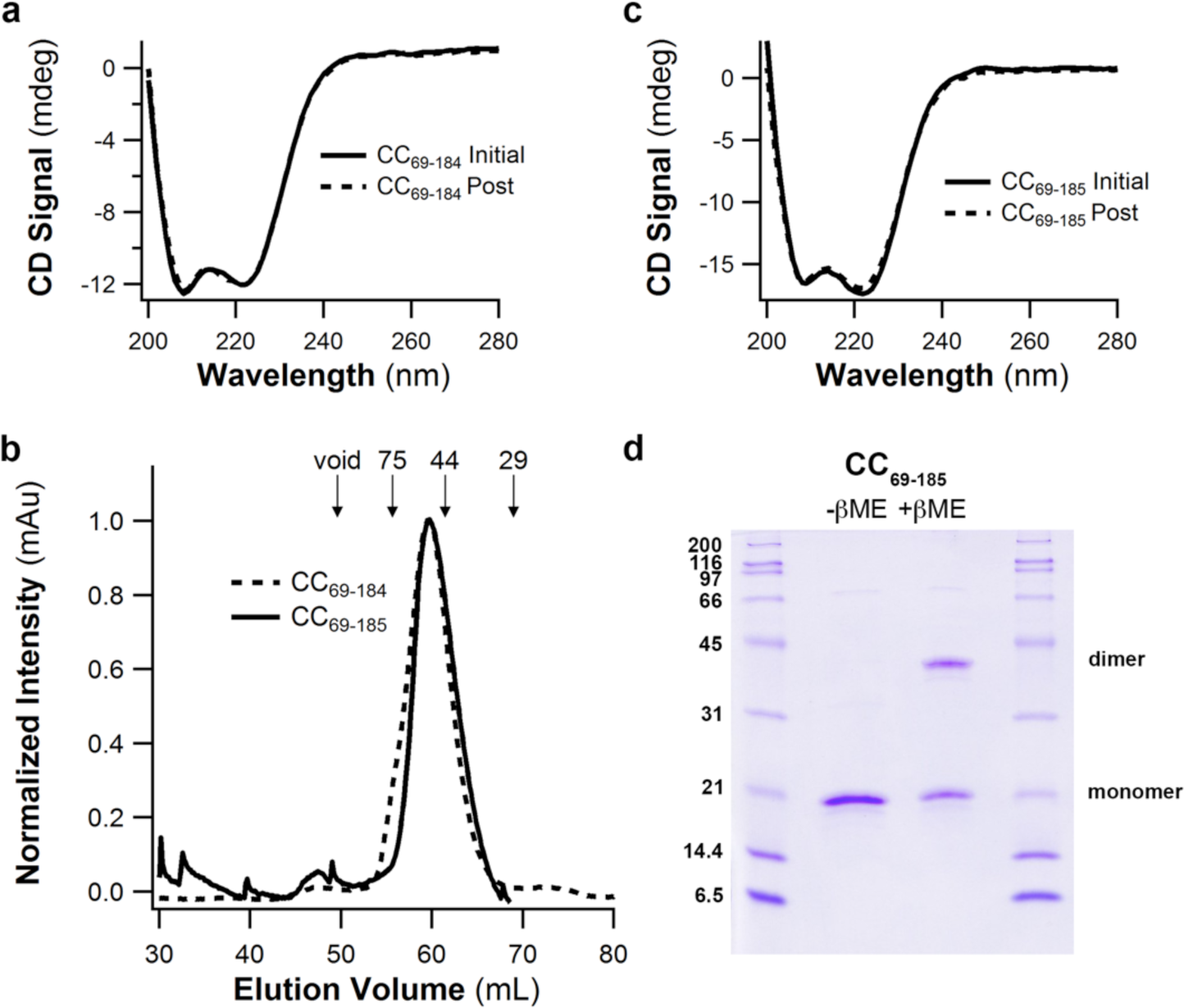
Biochemical characterization of CC_69-184_ and CC_69-185._ Related to Fig. 3. (a) Circular dichroism spectra of CC_69-184_ reveal alpha-helical signatures and reversible thermal unfolding (b) Superdex75 preparative SEC traces of CC_69-184_ and CC_69-185_ reveal a total molecular mass of 53 kDa based on a standard calibration curve. Given the calculated mass of 13.7 kDa/monomer the species is a tetramer. (b) Circular dichroism spectra of CC_69-185_ reveal an alpha-helical signature and reversible thermal unfolding. (d) SDS-PAGE analysis of CC_60-185_ under non-reducing conditions demonstrate predominantly disulfide-dependent dimer species.

**Supplemental Figure S4.**
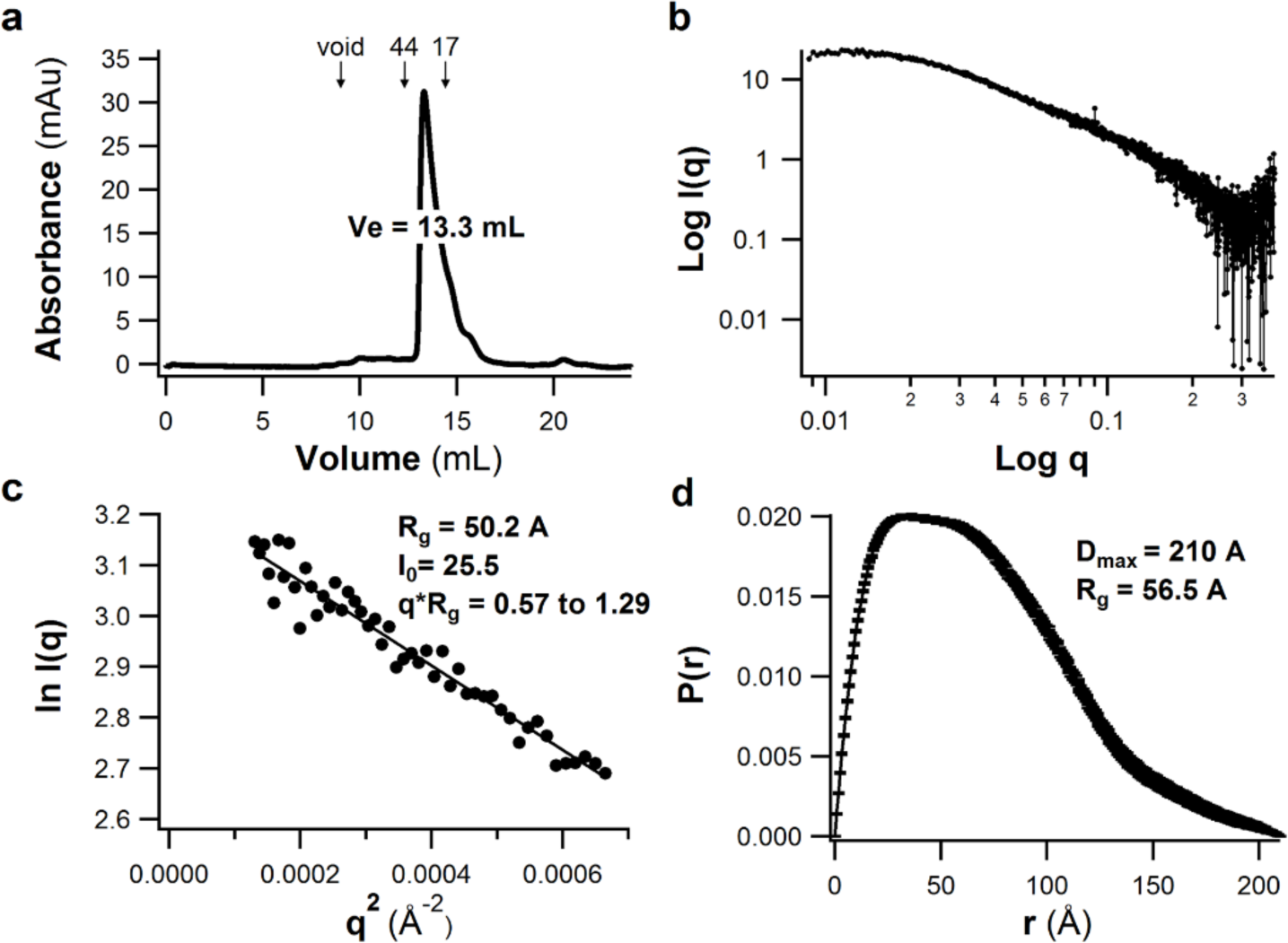
CC_69-185_ SEC-SAXS analysis. Related to Fig. 3. (a) Superdex-75 GL trace of CC_69-185_ obtained at the beamline reveals a monodisperse sample. (b) Scattering intensity profile. I(q): scattered intensity, q:scattering vector. (c) Guinier plot with calculated radius of gyration (Rg). (d) Pair wise distribution plot with calculated maximum particle size (Dmax) and radius of gyration (Rg).

**Supplemental Figure S5.**
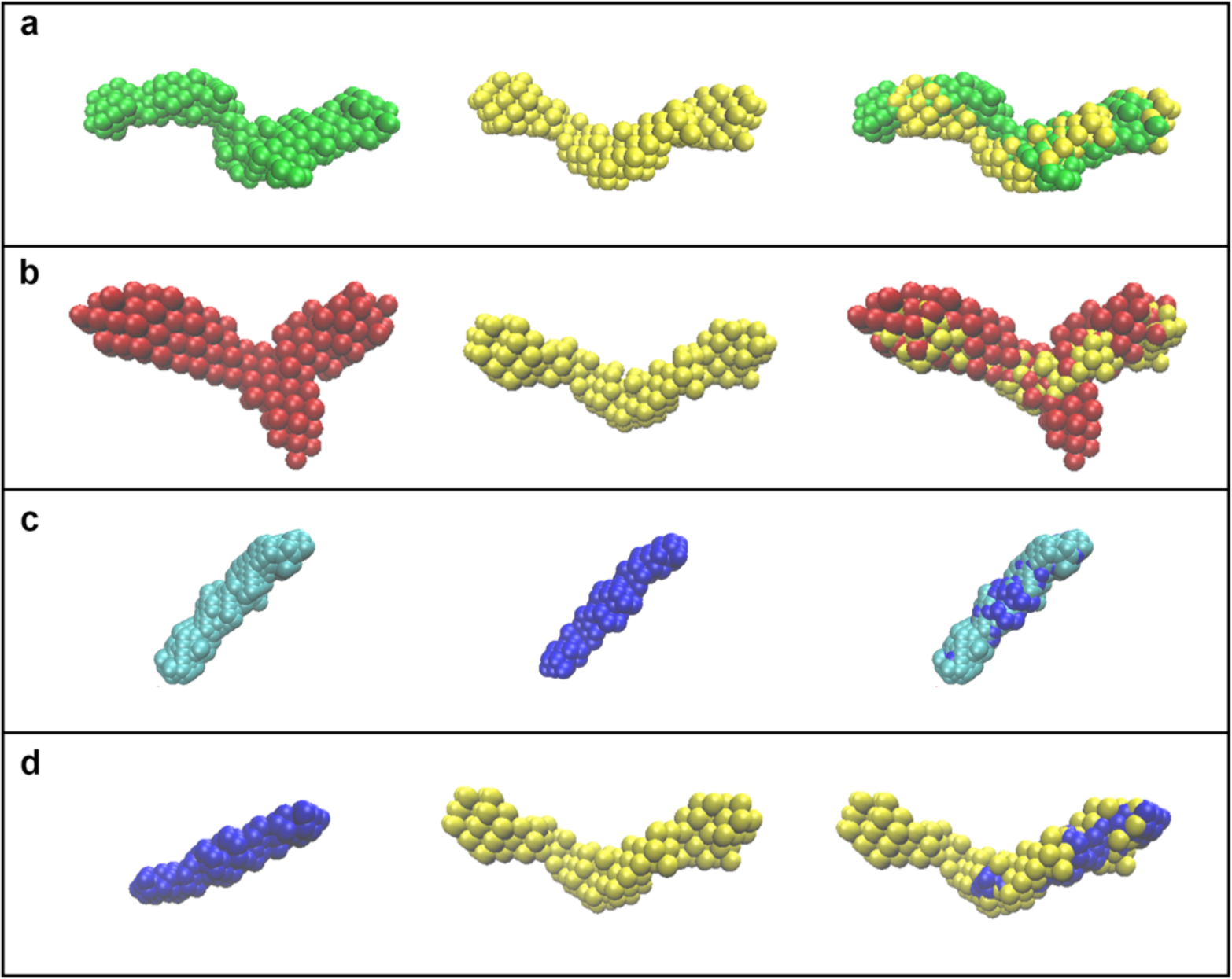
Pairwise comparisons of *ab initio* SEC-SAXS models. Related to Fig. 3 and Table 2. Molecular envelopes of (a) CC_69-185_ (b) CC_60-185_ (c) CC_112-185_ with no (P1) symmetry (left), P2 symmetry (middle) and superpositions with SUPCOMB (right). (d) Molecular envelopes of CC_112-185_ with P2 symmetry (left), CC_69-185_ with P2 symmetry (middle) and superposition by manual manipulation in PyMOL (right).

**Supplemental Figure S6.**
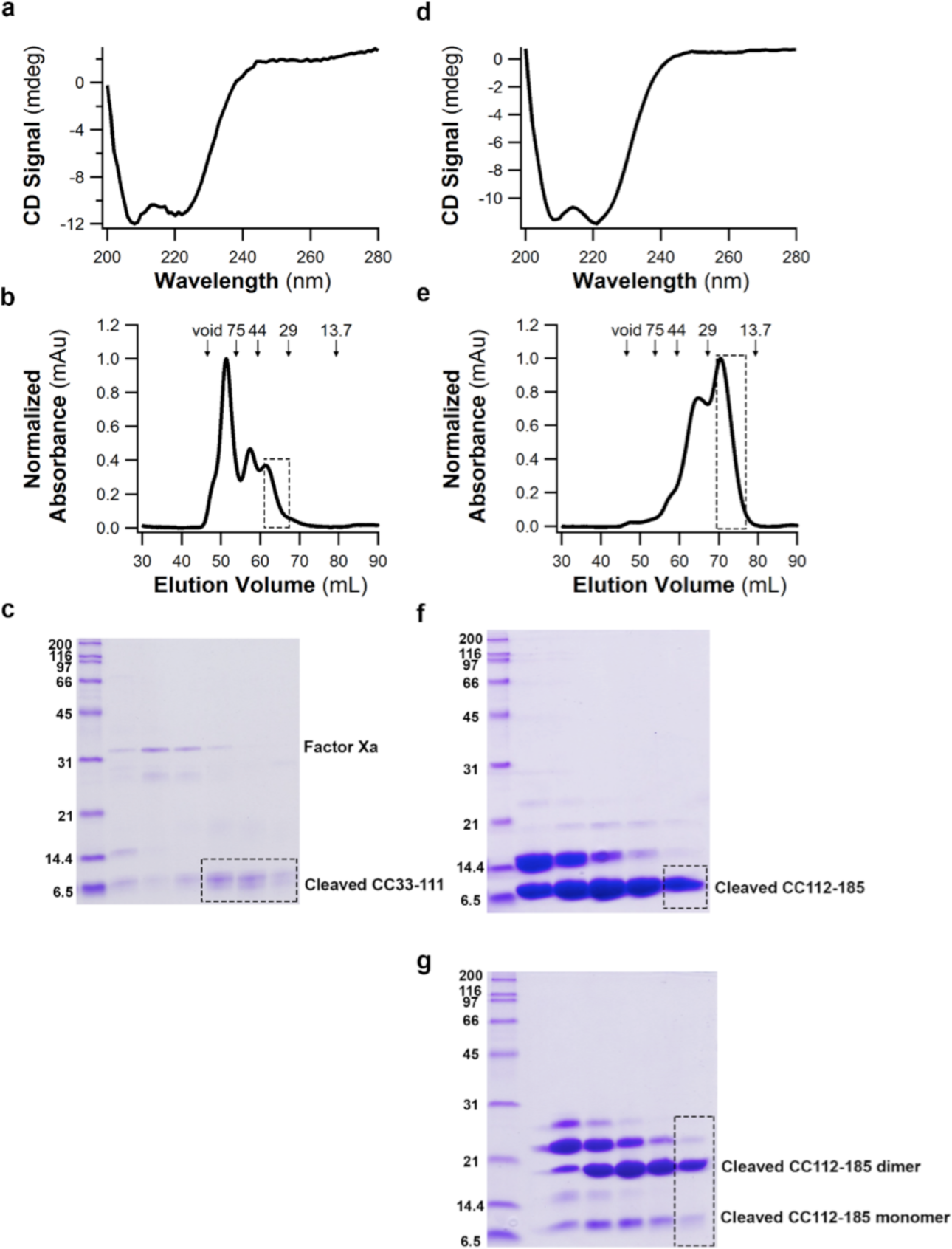
Biochemical characterization of CC_33-111_ and CC_112-185._ Related to Fig. 3. (a) Circular dichroism spectrum of CC_33-111_ reveals an alpha-helical signature. (b) Superdex-75 size exclusion chromatogram of CC_33-111._ The first peak contains uncleaved and cleaved CC_33-111,_ the second peak contains mostly Factor Xa protease, and the third peak (indicated by a dashed box) contains cleaved CC_33-111_ used in subsequent experiments. The molecular weight of the peak within the dashed box is 40 kDa based on a standard calibration curve, which consistent with a 4 or 5-mer of an ~8.8 kDa CC_33-111_ monomer. (c) SDS-PAGE analysis of CC_33-111_ fractions from Superdex- 75. Dashed box indicates fractions concentrated for SEC-SAXS. (d) Circular dichroism spectrum of CC_112-184_ reveals an alpha-helical signature. (e) Superdex-75 size exclusion chromatogram of CC_112-185._ The first peak contains uncleaved and cleaved CC_112-185,_ and the second peak (indicated by a dashed box) contains mostly cleaved CC_112-185_ used in subsequent experiments. The molecular mass of the species is 23 kDa based on a standard calibration curve and thus consistent with a 2- or 3-mer of an ~9 kDa CC112-185 species. (f) SDS-PAGE analysis of CC_112-185_-containing fractions from Superdex 75 showin in (e). Dashed box indicates fraction selected for SEC-SAXS.

**Supplemental Figure S7.**
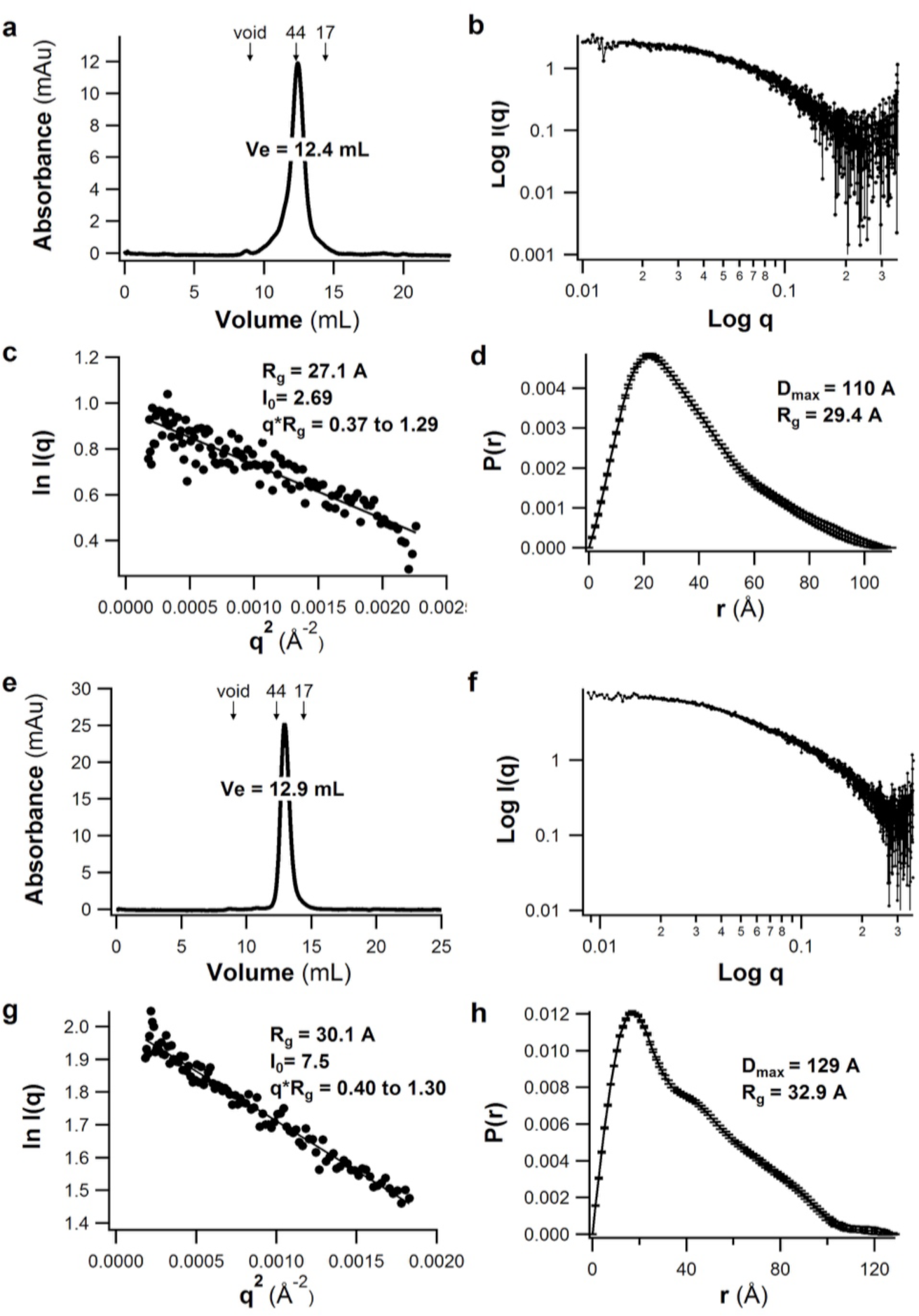
CC_33-111_ and CC_112-185_ SEC-SAXS analysis. Related to Fig. 3. (a) Superdex-75 GL trace of CC_33-111_ obtained at the beamline reveals monodisperse sample with a molecular weight of 44 kDa. Given the monomeric molecular mass of ~8.8 kDa the species is an apparent 5-mer. (b) Scattering intensity profile. I(q): scattered intensity, q:scattering vector. (c) Guinier plot with calculated radius of gyration (Rg). (d) Pair wise distribution plot with calculated maximum particle size (Dmax) and radius of gyration (Rg). (e) Superdex-75 GL trace of CC_112-185_ obtained at the beamline reveals monodisperse sample ( f) Scattering intensity profile. I(q): scattered intensity, q:scattering vector. (g) Guinier plot with calculated radius of gyration (Rg). (h) Pair wise distribution plot with calculated maximum particle size (Dmax) and radius of gyration (Rg).

**Supplemental Figure S8.**
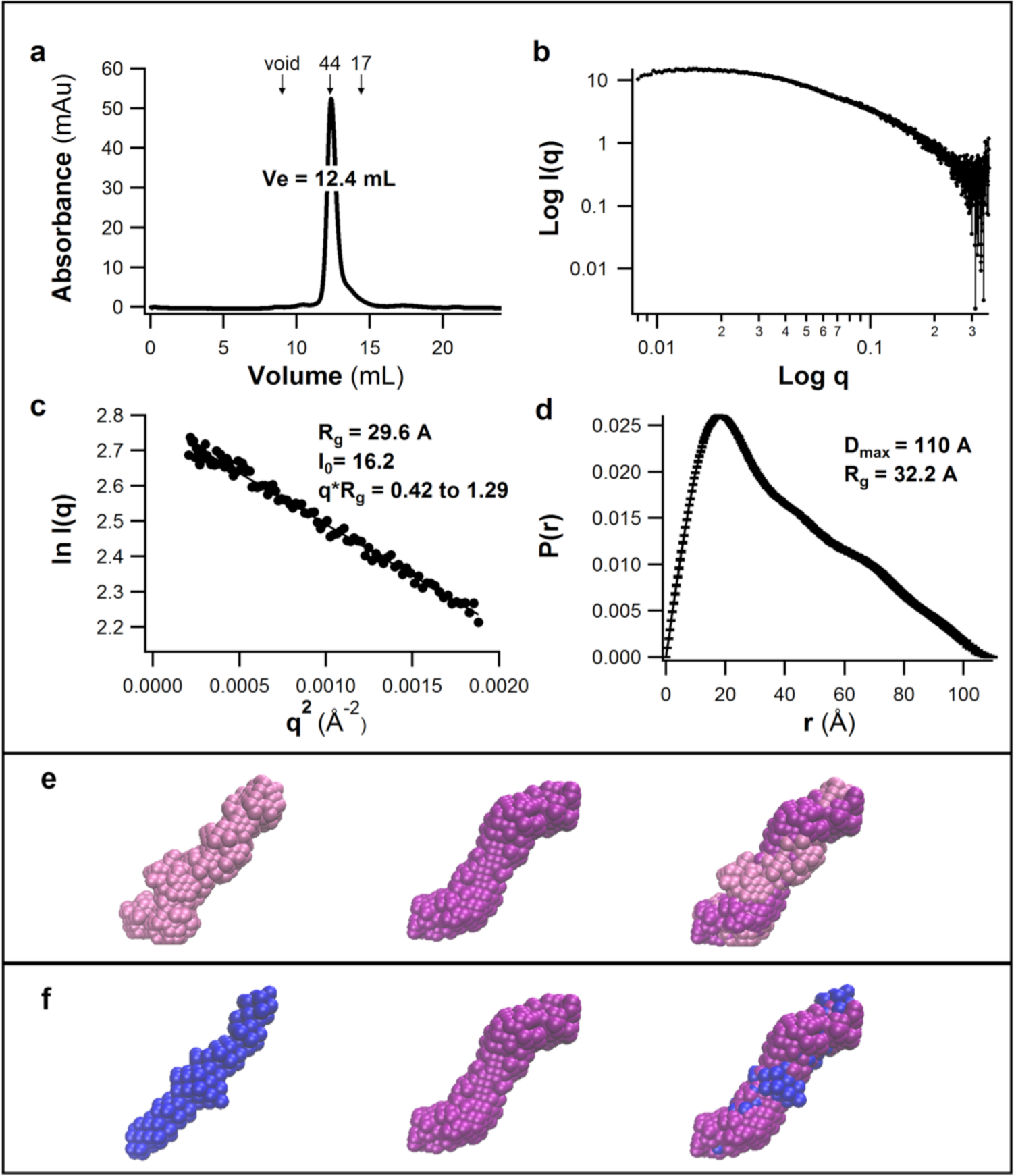
mLZ_93-171_ SEC-SAXS analysis. Related to Fig. 3. (a) Superdex-75 GL trace of mCC_93-171_ reveals monodisperse sample (b) Scattering intensity profile. I(q): scattered intensity, q:scattering vector. (c) Guinier plot with calculated radius of gyration (Rg). (d) Pair wise distribution plot with calculated maximum particle size (Dmax) and radius of gyration (Rg). (e) Molecular envelopes of mCC_93-171_ with no (P1) symmetry constraints (left), P2 symmetry constraints (middle), and the two envelopes superimposed with SUPCOMB (right). (f) Molecular envelopes of CC_112-185_ with P2 symmetry constraint (left), mCC_93-171_ with P2 symmetry constraint (middle) and superposition with SUPCOMB (right).

**Supplemental Figure S9.**
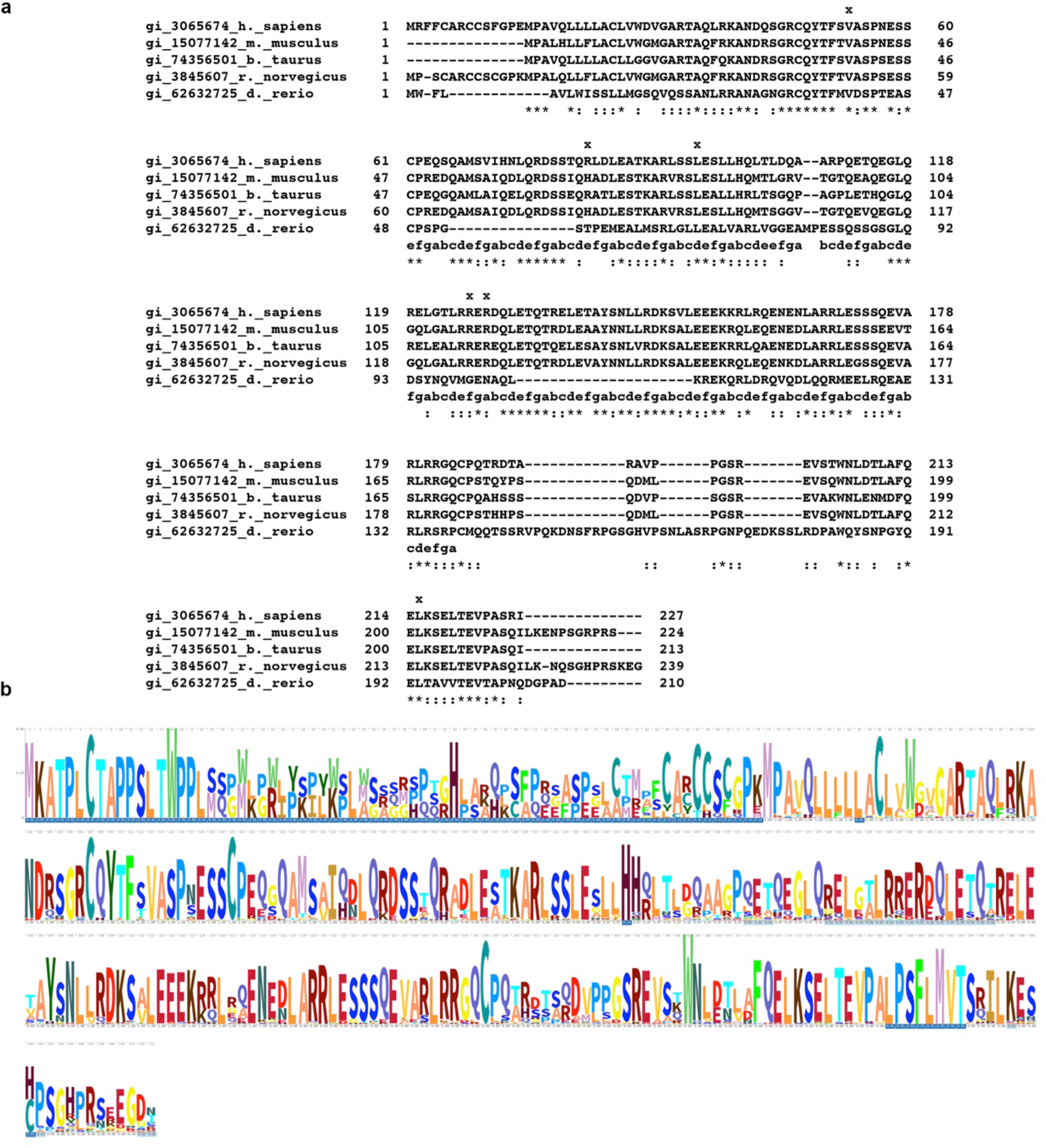
Related to Fig. 1. Sequence comparison of myocilins across organisms. (a) Multiple sequence alignment of N-terminal region of myocilin from human, mouse, bovine, rat, and zebrafish (OLF domain excluded). (b) HMM weblogo representation of conservation across the same region as in (a) across myocilin from 75 species.

**Supplemental Figure S10.**
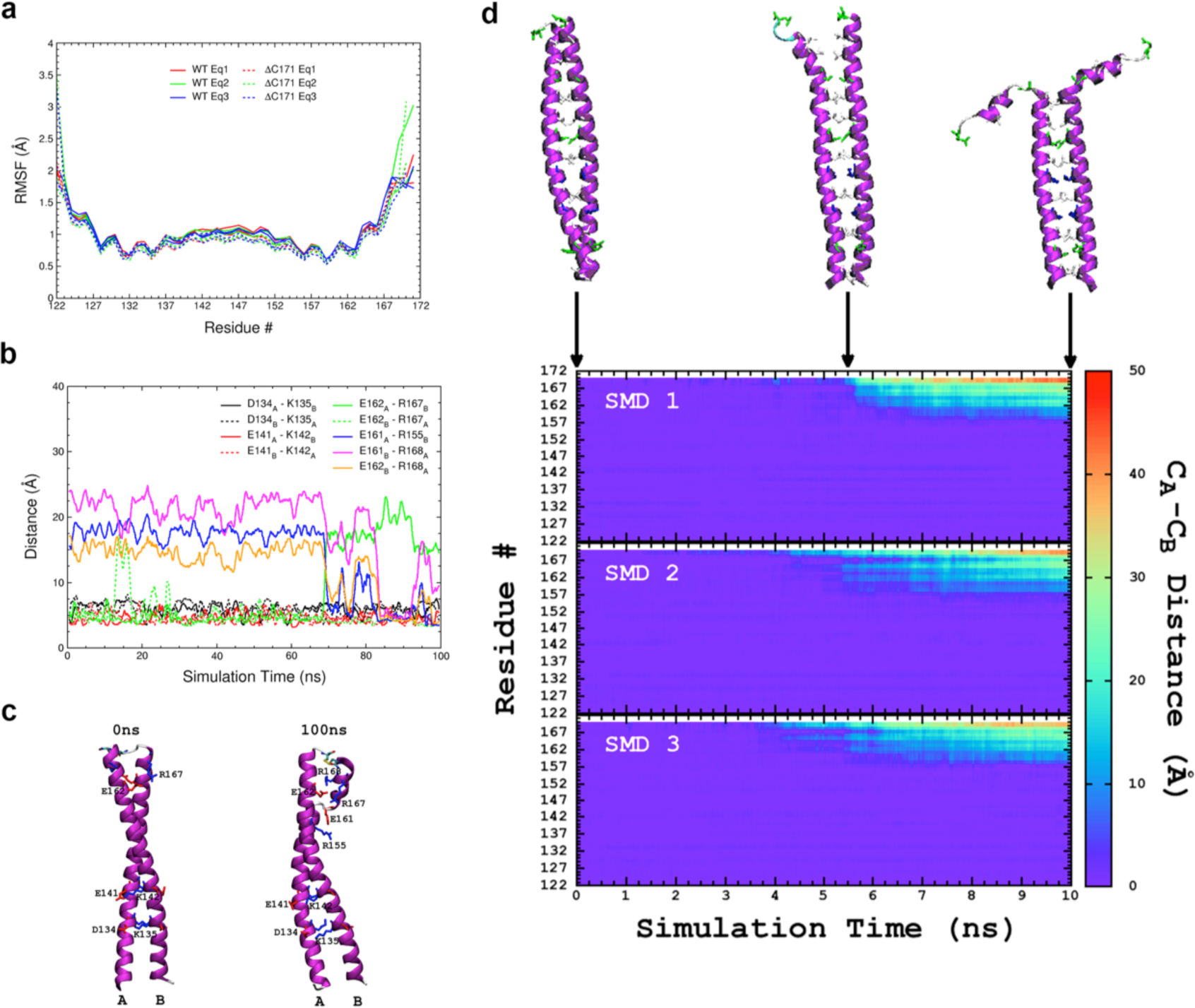
Molecular dynamics simulations of mLZ_122-171._ Related to Fig. 4. (a) Root mean squared fluctuation (RMSF) values for individual mLZ_122-171_ residues for simulations conducted at 310 K. (b)/(c) Salt bridges formed between individual coils A and B during 100-ns simulation of mLZ_122-171_ at 400 K. For (c), backbone structures are shown for 0 ns (left) and 100 ns (right) of simulation time with salt-bridge-forming residues shown explicitly. Salt bridges were identified by ≤3.2 Å distance between side chain oxygen and nitrogen atoms. (d) Distances between backbone carbonyl carbon atoms of chains A and B for each residue of mLZ_122-171_ during steered molecular dynamics (SMD) simulations at 310 K. Backbone structures from the SMD1 trajectory are shown above the graph with side chains of heptad positions ‘*a*’ and ‘*d*’ shown explicitly. For all protein structures shown: Individual protein chains are labeled A and B. Protein backbone is shown in cartoon representation colored by secondary structure: (magenta) α-helix and (white) random coil. Explicit side chains are shown in licorice representation colored by residue type: (blue) positively charged, (red) negatively charged, (green) polar, and (white) hydrophobic.

**Supplemental Figure S11.**
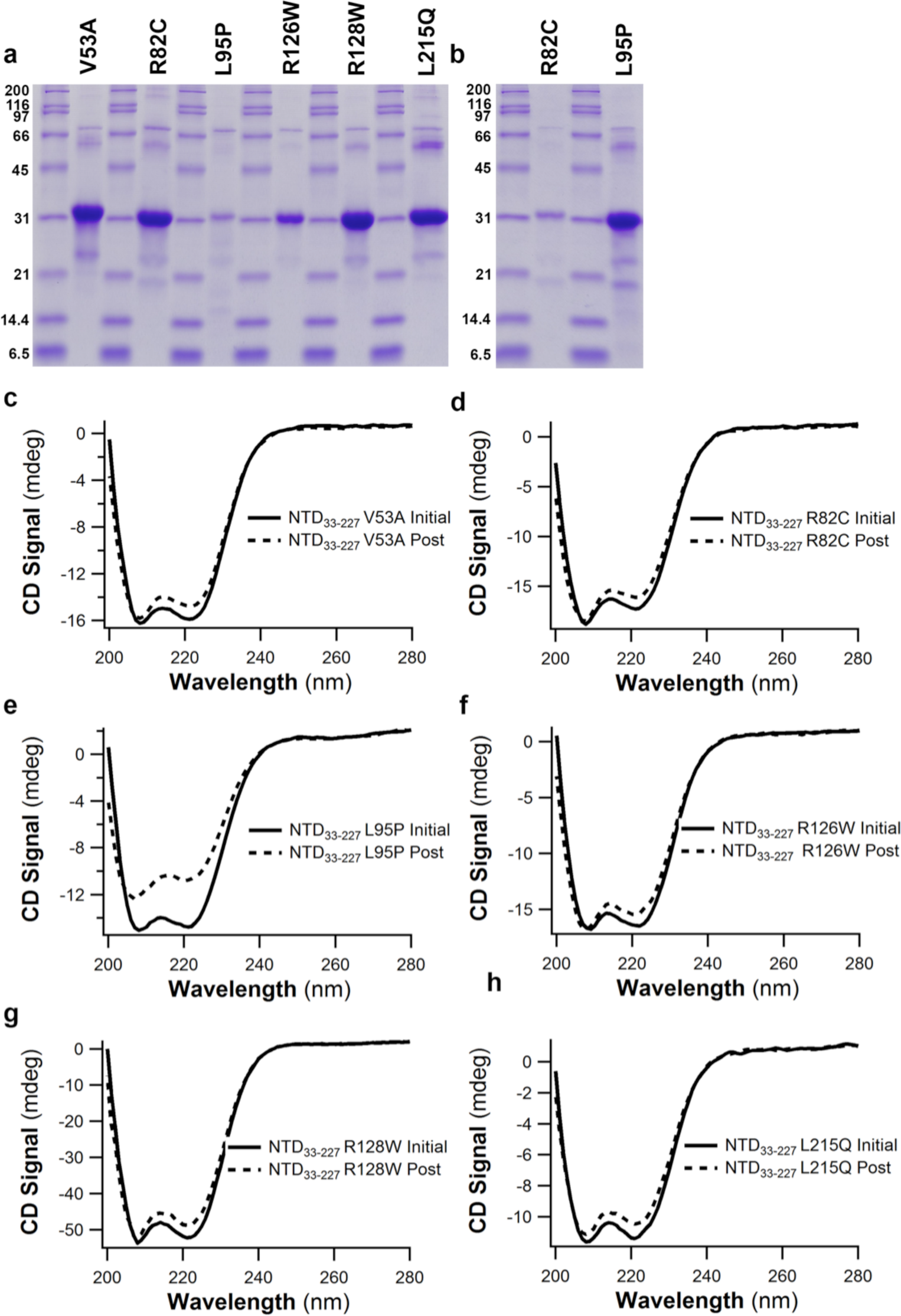
Biochemical characterization of glaucoma-associated NTD33-227 variants. Related to Fig. 4. (a) SDS-PAGE analysis of purified variants (~ 30 kDa monomer). (bh) Comparison of CD spectra before (initial) and after (post) thermal melt shows a high level of reversibility for six variants studied.

**Supplemental Figure S12.**
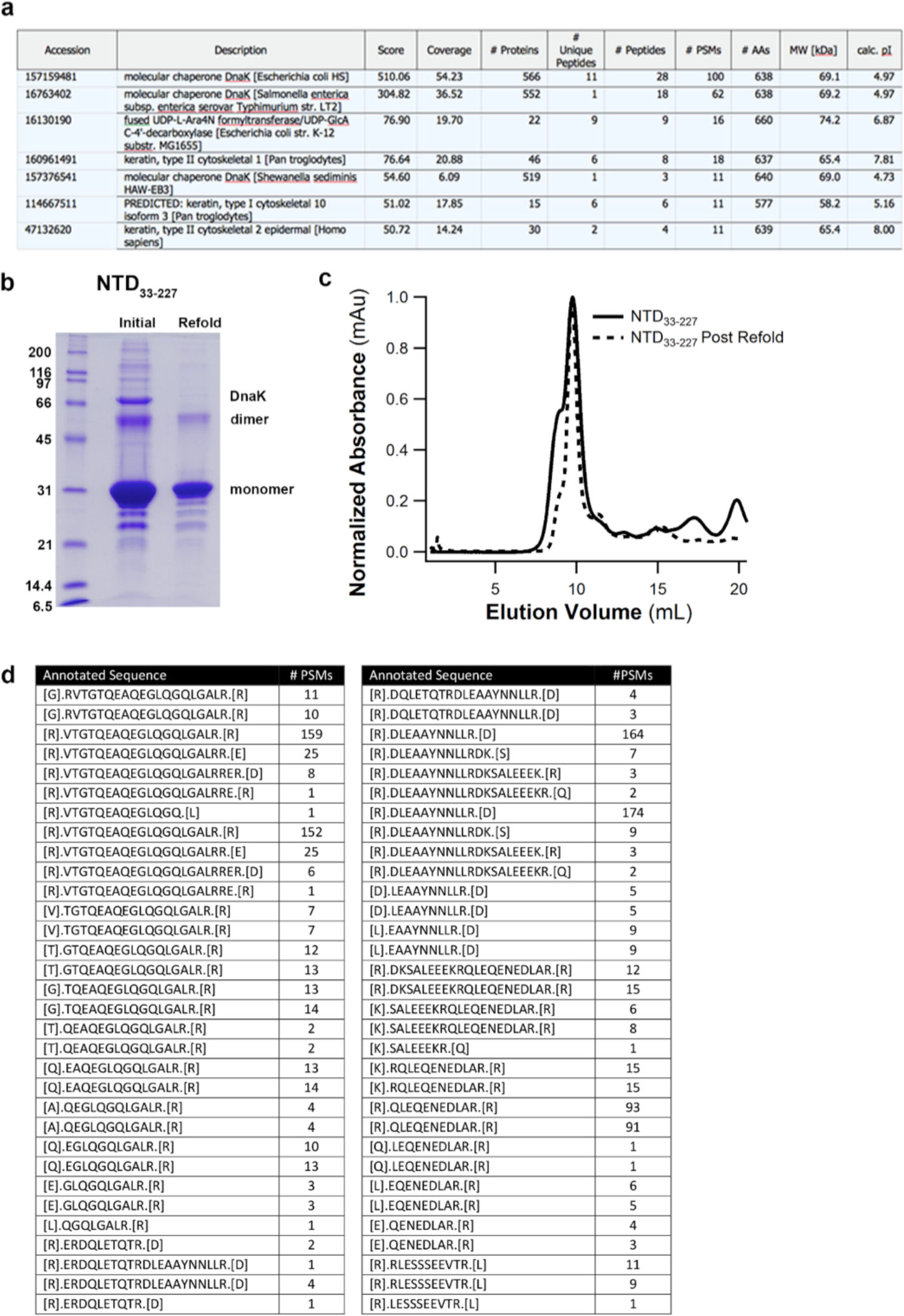
Additional supplemental material accompanying methods. Related to Supplemental Methods. (a) Identification of DnaK by mass spectrometry as main contaminant (~60 kDa) visible after purification on SDS-PAGE. (b) SDS-PAGE analysis of NTD_33-227_ before and after unfolding/refolding procedure. (c) Comparison of Superdex-75 elution profile of two samples in (b). (d) Identification of mLZ_93-171_ as predominant product after cleavage of LZ_55-171_ by Factor Xa.

**Supplemental Table S1.**
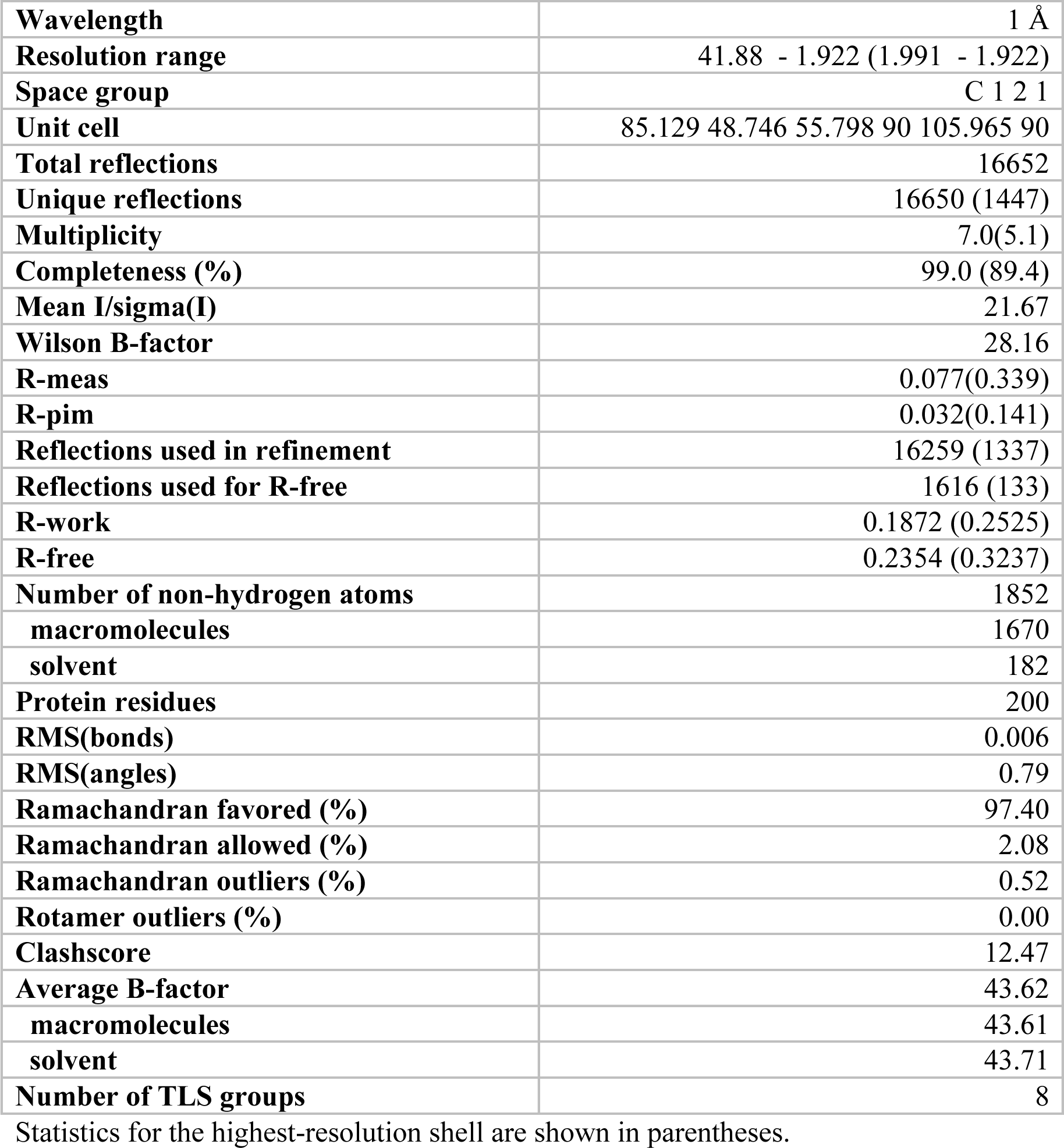
Related to Fig. 4. Crystallographic statistics.

|  |  |
| --- | --- |
| **Wavelength** | 1 Å |
| **Resolution range** | 41.88 - 1.922 (1.991 - 1.922) |
| **Space group** | C 1 2 1 |
| **Unit cell** | 85.129 48.746 55.798 90 105.965 90 |
| **Total reflections** | 16652 |
| **Unique reflections** | 16650 (1447) |
| **Multiplicity** | 7.0(5.1) |
| **Completeness (%)** | 99.0 (89.4) |
| **Mean I/sigma(I)** | 21.67 |
| **Wilson B-factor** | 28.16 |
| **R-meas** | 0.077(0.339) |
| **R-pim** | 0.032(0.141) |
| **Reflections used in refinement** | 16259 (1337) |
| **Reflections used for R-free** | 1616 (133) |
| **R-work** | 0.1872 (0.2525) |
| **R-free** | 0.2354 (0.3237) |
| **Number of non-hydrogen atoms** | 1852 |
| **macromolecules** | 1670 |
| **solvent** | 182 |
| **Protein residues** | 200 |
| **RMS(bonds)** | 0.006 |
| **RMS(angles)** | 0.79 |
| **Ramachandran favored (%)** | 97.40 |
| **Ramachandran allowed (%)** | 2.08 |
| **Ramachandran outliers (%)** | 0.52 |
| **Rotamer outliers (%)** | 0.00 |
| **Clashscore** | 12.47 |
| **Average B-factor** | 43.62 |
| **macromolecules** | 43.61 |
| **solvent** | 43.71 |
| **Number of TLS groups** | 8 |

Statistics for the highest-resolution shell are shown in parentheses.

**Supplemental Table S2.**
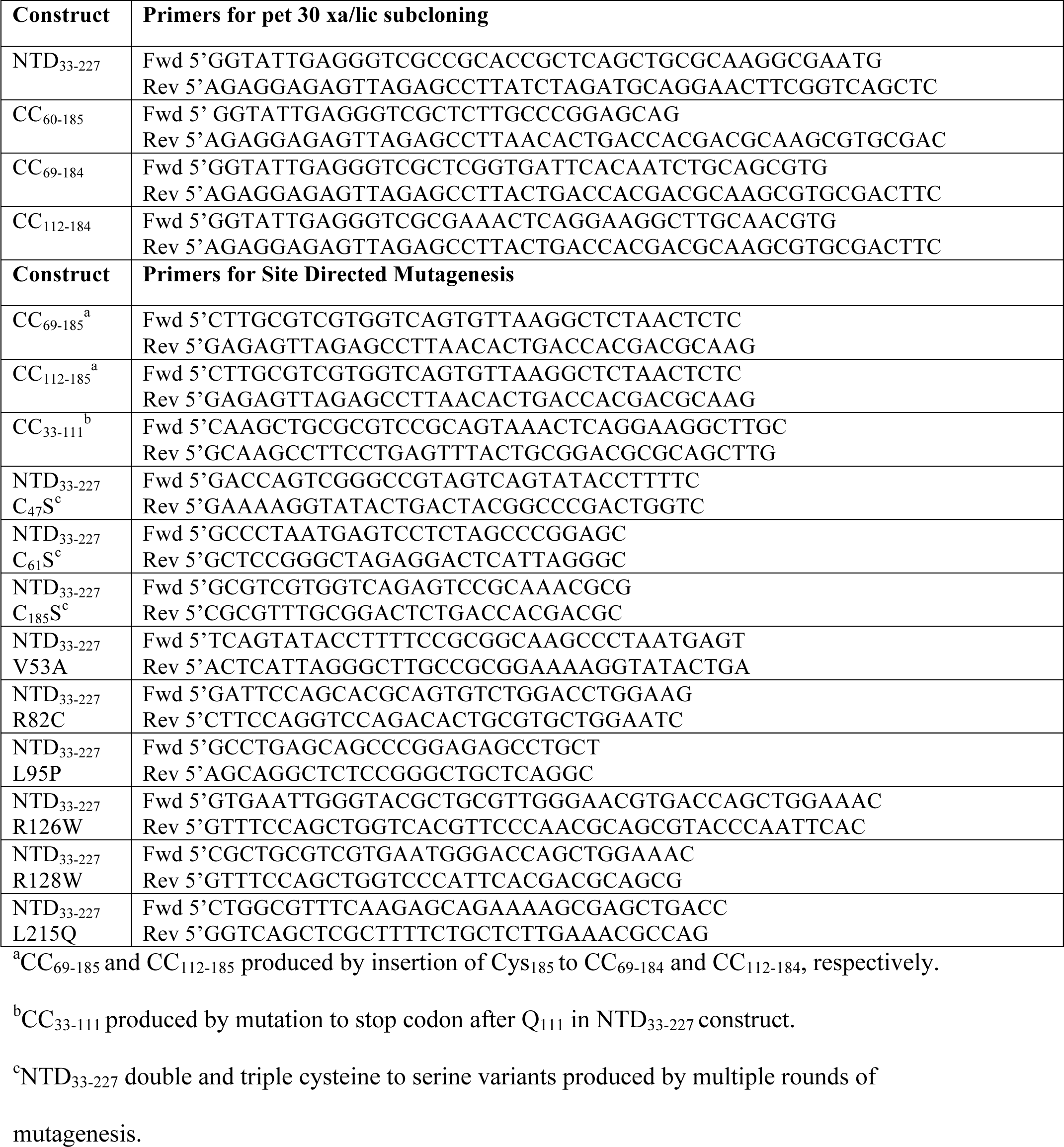
Related to Supplemental Methods. Primers used in this study.

